# Single cell profiling of total RNA using Smart-seq-total

**DOI:** 10.1101/2020.06.02.131060

**Authors:** Alina Isakova, Norma Neff, Stephen R. Quake

**Affiliations:** Department of Bioengineering, Stanford University, Stanford, California, USA; Department of Applied Physics, Stanford University, Stanford, California, USA; Chan Zuckerberg Biohub, San Francisco, California, USA

## Abstract

The ability to interrogate total RNA content of single cells would enable better mapping of the transcriptional logic behind emerging cell types and states. However, current RNA-seq methods are unable to simultaneously monitor both short and long, poly(A)+ and poly(A)-transcripts at the single-cell level, and thus deliver only a partial snapshot of the cellular RNAome. Here, we describe Smart-seq-total, a method capable of assaying a broad spectrum of coding and non-coding RNA from a single cell. Built upon the template-switch mechanism, Smart-seq-total bears the key feature of its predecessor, Smart-seq2, namely, the ability to capture full-length transcripts with high yield and quality. It also outperforms current poly(A)–independent total RNA-seq protocols by capturing transcripts of a broad size range, thus, allowing us to simultaneously analyze protein-coding, long non-coding, microRNA and other non-coding RNA transcripts from single cells. We used Smart-seq-total to analyze the total RNAome of human primary fibroblasts, HEK293T and MCF7 cells as well as that of induced murine embryonic stem cells differentiated into embryoid bodies. We show that simultaneous measurement of non-coding RNA and mRNA from the same cell enables elucidation of new roles of non-coding RNA throughout essential processes such as cell cycle or lineage commitment. Moreover, we show that cell types can be distinguished based on the abundance of non-coding transcripts alone.

## MAIN

Efforts in characterizing transcriptional states of single cells have so far mostly focused on protein-coding RNA^1–4^. However, a growing number of studies indicate that non-coding RNAs (ncRNAs), are actively involved in cell function and specialization^5–8^. Importantly, compared to the coding RNA, which is transcribed from only ~1-2% of the genome, the non-coding RNA constitutes a major fraction of all cellular transcripts and covers ~70% of the genomic content^9^. The role of these transcripts in shaping different cell types and states remains poorly understood.

Several groups have demonstrated the possibility of measuring the levels of ncRNA in single cells^10,11^. The respective methods, however, are designed to target only a subset of non-coding transcripts, which are either short (~18-200 nt, e.g. microRNA)^11,12^ or long (>200 nt, e.g. lncRNA or circRNA)^10,13–15^, while none of them offer a simultaneous assessment of all RNA types within a cell. This limits one’s ability to map the regulatory connection between coding, and different types of non-coding transcripts within a cell and calls for the development of novel single-cell technologies capable of assaying both poly(A)^+^ and poly(A)^-^ RNA, irrespective of transcript length.

In the present study we describe Smart-seq-total, a scalable method designed to capture both coding and non-coding transcripts regardless of their length. Inspired by the widely used Smart-seq2 protocol^16^, this method harnesses template switching capability of MMLV reverse transcriptase to generate full-length cDNA with high yield and quality. In addition, Smart-seq-total is designed to capture non-polyadenylated RNA through template-independent addition of polyA tails and further oligo-dT priming of all cellular transcripts. Therefore, Smart-seq-total simultaneously measures cellular levels of mRNA alongside other RNA types in the same cell, which permits the discovery of non-coding regulatory patterns of a cell and at the same time facilitates the integration of this data with the existent single cell RNA-seq datasets.

Smart-seq-total relies on the ability of *E.coli* poly(A) polymerase to add adenine tails to the 3’ prime of RNA molecules. Total polyadenylated RNA is then reverse transcribed using anchored oligo dT, in the presence of the template switch oligo (TSO)^17^ (**Fig. 1a**). Compared to previous studies that explored similar approaches to construct libraries from total RNAs^18,19^, Smart-seq-total utilizes an optimized version of the (TSO^16^), specifically engineered to be rapidly eliminated from the reaction directly following the reverse transcription. This allows us to remove the “contaminant” constructs, originating from polyA-tailing and mispriming of TSO, which otherwise dominate the resulting sequencing library and render the short RNA transcripts undetectable (see **Supplementary Fig. 1a-b**). Additionally, we employ a CRISPR-mediated removal of overrepresented sequences, which allows us to eliminate the majority of the sequences corresponding to ribosomal RNA from the final library in a single-pool reaction (targeting 69 rRNA regions**; see Supplementary table 1**). Applied to single HEK293T cells Smart-seq-total identified, alongside mRNA, a broad spectrum of non-coding RNA genes, such as snoRNA, scaRNA, histone RNA and lncRNA. The majority of these molecules endogenously lack poly(A) tails and thus cannot be captured through a direct polyA-priming employed by Smart-seq2 or other popular scRNA-seq methods^1^ (**Fig. 1b; see Supplementary Fig. 1c**). Among other ncRNA, detected uniquely by Smart-seq-total are tRNAs and mature miRNAs (**Fig. 1c; see Supplementary Fig. 1d**).

**Figure 1.**
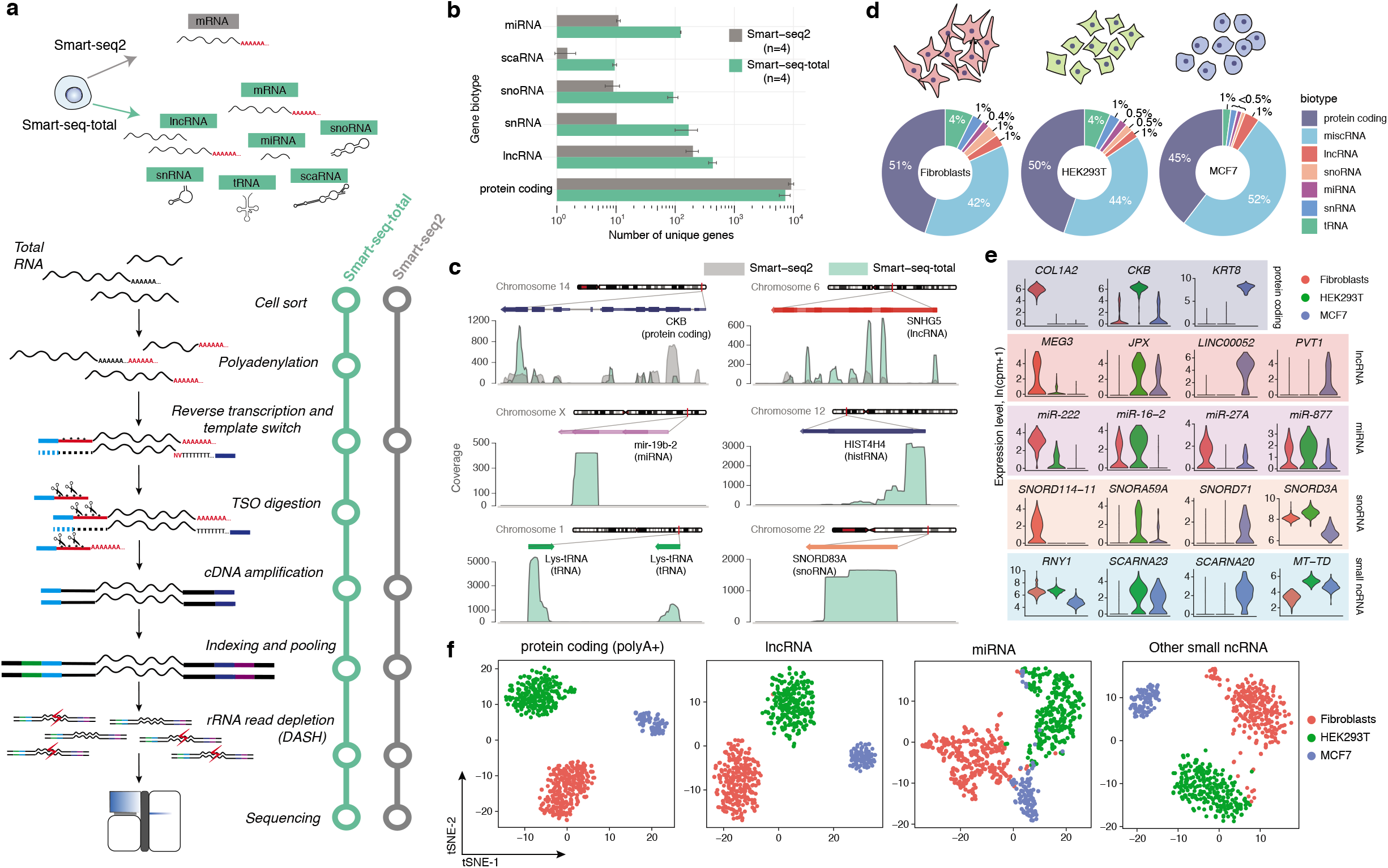
Smart-seq-total performance. **a. Schematic comparison of Smart-seq2 and Smart-seq-total pipelines.** Following cell lysis, total cellular RNA is polyadenylated, primed with anchored oligodT and reverse transcribed in a presence of the custom degradable TSO. After reverse transcription, TSO is enzymatically cleaved, single-stranded cDNA is amplified and cleaned up. Amplified cDNA is then indexed, pooled and depleted from ribosomal sequences using DASH^33^. Resulting indexed libraries are then pooled and sequenced on Illumina platform. **b. Average number of genes per biotype detected by Smart-seq2 and Smart-seq-total in single HEK293T cells.** Genes were assigned to a specific biotype based on GENCODE v32 annotation for the reference genome. tRNA was quantified using high-confidence gene set obtained from GtRNAdb. Adaptor-depleted libraries, for both Smart-seq2 and Smart-seq-total, were depth normalized to ~2.5 Mio reads per cell. Error bars denote standard deviation (n=4). **c. Coverage of multiple gene biotypes shown for Smart-seq2 and Smart-seq-total data.** Computed as a sum of n=4 cells. **d. Distribution of mapped reads across RNA types in human primary fibroblasts, HEK293T and MCF7 cells.** Percentage of total reads mapped to each RNA type. **e. Examples of coding and non-coding marker genes for each cell type.** Top exemplary markers per biotype computed among cell types using Wilcoxon Rank Sum test. *RNY1* belongs to miscRNA, *SCARNA23* and *SCARNA20* – to scaRNA, *MT-TD* – to mitochondrial tRNA class. **f. t-SNE plots of three profiled human cell types generated using indicated subset of genes.** From left to right: protein coding, lncRNA, miRNA and other small ncRNA (include snoRNA, snRNA, scaRNA, scRNA and miscRNA). We have excluded histone coding genes from protein coding (polyA+) set, since a large fraction of these RNAs are known to lack polyA tails^34^.

To assess the scalability of the method, we sequenced total RNA from individual human primary dermal fibroblasts (n=277), HEK293T (n=245) and MCF7 (n=90) cells sorted in 384-well plates and processed in 1/10 of the standard Smart-seq2 volume^16^. Within all three cell types we identified a broad spectrum of transcripts such as mRNA, miRNA, lncRNA, and snoRNA in each profiled cell (**Fig. 1d; see Supplementary Fig. 2a-c**). We found metazoan cytoplasmic RNA7SK and RN7SL1, annotated as ‘miscellaneous RNA’ type (miscRNA) in GENCODE database, to be the most abundant in our data comprising together ~40 % of all mapped reads (**Fig. 1d; see Supplementary Fig. 3**). Among cell-type specific transcripts we found well-characterized marker genes for either fibroblasts (*COL1A2, FN1, MEG3*), HEK293T (*CKB, AMOT, HEY1*) or MCF7 cells (*KRT8, TFF1*) (**Fig. 1e; see Supplementary Fig. 4a**) as well as transcripts which belong to various types of ncRNA, such as microRNA, snoRNA and lncRNA (**Fig. 1e; see Supplementary Fig. 4**). For example, we found high levels of *MIR222* in fibroblasts while could not detect it in MCF7 cells. We also observed that oncogenic miRNA cluster *MIR17HG* is specific to HEK293T cells, while not found in neither fibroblasts nor MCF7 cells. In contrast, MCF7-specific transcripts include lncRNA, such as *LINC00052*, as well as snoRNA, such as *SNORD71* and *SNORD104.*

Given the observed differences in the levels of non-coding RNA across profiled cells, we next asked whether non-coding RNA alone could be used to distinguish cell types. To answer this question, we performed principal component analysis (PCA) followed by the dimensionality reduction on the genes corresponding to one or multiple ncRNA types. Evaluation of the similarity between cells in two-dimensional space revealed that, taken alone, lncRNA, and miRNA separate the investigated cell types into three distinct clusters, while combining snoRNA, scaRNA, snRNA and tRNA together allowed us to achieve similar results (**Fig. 1f**).

In addition to cell-type dependent differences in ncRNA, the abundance of certain non-coding transcripts also changed throughout the cell cycle (**Fig. 2a**). In agreement with previous bulk studies, suggesting the involvement or miRNA in cell-cycle regulation^20,21^, we found that levels of a subset of miRNAs in a cell dynamically change through the cell cycle, peaking at either S, G2M or G1 phase (**Fig. 2a**). For example, our data showed that the levels of *MIR16-2* in fibroblasts are high during the S phase and later gradually decrease during G2M and G1 phases **(see Supplementary Fig. 5**). The opposite holds true for *MIR222*, in both fibroblasts and HEK293T cells, which is upregulated during cell proliferation (G1) and decays during DNA replication (S) and cell division (G2M) phases (**Fig. 2a; see Supplementary Fig. 6**). Among miRNAs upregulated during G2M phase we identified *MIR27A, MIR103A2*, and *MIR877* (**see Supplementary Fig. 5-7**). In addition to miRNA, a large number of lncRNA, snRNA, scaRNA snoRNA and miscRNA were also upregulated during the G2M phase (**Fig. 2a, see Supplementary Fig. 5-7**). Given the active role of these RNA types in splicing and ribosome biogenesis, we suggest that they are produced by the cell in response to a rapid demand for protein synthesis and cell growth during the G2M phase.

**Figure 2.**
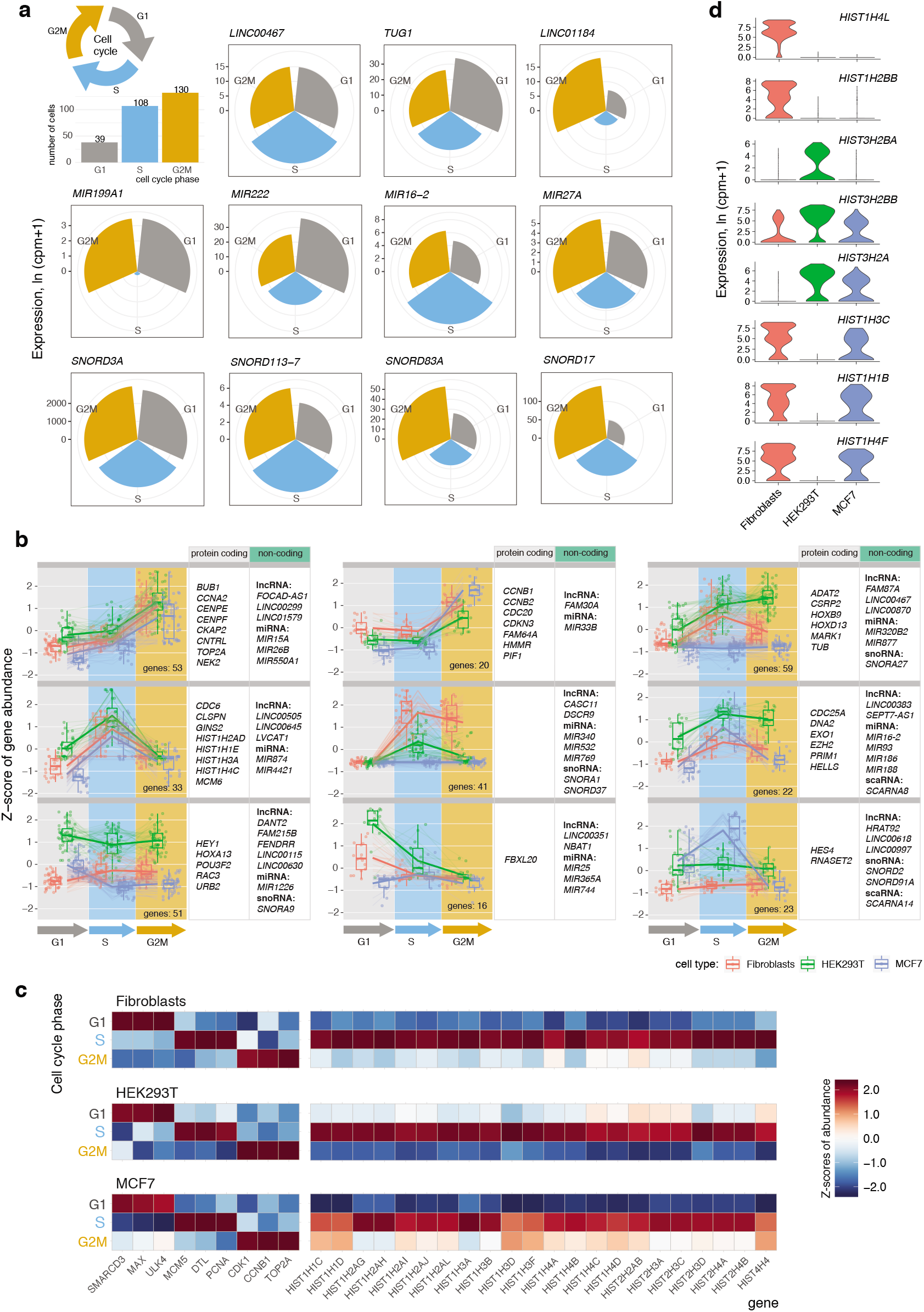
Dynamics of cellular non-coding transcripts throughout the cell cycle. **a. Cell-cycle dependent expression of non-coding genes.** Examples of lncRNA, miRNA and snoRNA differentially expressed throughout the cell cycle in human primary dermal fibroblasts. Circular charts depict average expression of a given gene across all cells identified to be in a certain phase of the cell cycle. **b. Cell-cycle specific gene clusters comprised of coding and non-coding RNA.** Clusters were identified through hierarchical clustering of top 750 mRNA differentially expressed during the cell-cycle and all non-coding genes expressed in at least one phase. **c. Expression of known cell-cycle and histone genes across G1, S and G2M phases.** A curated list of histone RNA detected in all three cell types is shown. **d. Examples of histone mRNA differentially expressed between three profiled cell types.** Top three marker histone genes per cell type are shown.

To further link the observed non-coding RNA dynamics with the expression of well-characterized cell-cycle mRNA markers, we searched for co-regulated coding and non-coding genes throughout the cell cycle. We identified 24 clusters comprised of co-expressed coding and non-coding genes specific to either one or multiple cell types (**Fig. 2b, see Supplementary Fig. 8, Supplementary Table 2**). Two of these mixed-gene clusters (33 genes upregulated in the S phase and 53 genes upregulated in the G2M phase) showed identical patterns in all three profiled cell types. Interestingly, both clusters are marked by landmark cellcycle genes, such as *CCNA2, MCM6* and *TOP2A*, but also include miRNAs, lncRNAs and snRNAs previously unknown to follow a distinct expression pattern upon transition between phases.

Histone RNA is another type of mainly non-polyadenylated RNA which we observed to be strongly correlated with the cell cycle. Consistent with prior studies^22,23^, histone RNA levels sharply rise during the S phase in all three profiled cell types (**Fig. 2c**). The ability to capture non-polyadenylated histones also has a strong impact on cell clustering, by introducing a cell cycle bias. Particularly, histones drive the separation of each cell type into two distinct populations (**see Supplementary Fig. 9a**), marked by increased levels of the majority of histone genes during the DNA replication phase (**see Supplementary Fig. 9b**).

In addition to being expressed in a cell cycle-dependent manner, we also identified several histones to be cell type specific. For example, *HIST1H4L* is expressed in fibroblasts but absent in HEK293T and MCF7 cells, while *HIST1H1B*, is absent in HEK293T cells while present in the other two cell types (**Fig. 2d**). Given the importance of histones in establishing and maintaining a distinct chromatin landscape of a cell, we anticipate that the ability to measure corresponding transcripts could be valuable for predicting the epigenetic state of a cell.

Finally, we sought to understand whether the unique non-coding signature acquired by different cell types is established during early stages of cell development and if so, how dynamic it is with respect to cellular transcriptome. To address this question we referred to an *in vitro* model of early lineage commitment: the differentiation of pluripotent stem cells into embryoid bodies^24^. The role of ncRNA in maintaining stem cell pluripotency and lineage commitment has been demonstrated previously through bulk experiments^25,26^. Thus, we hypothesized that applying Smart-seq-total to single cells at different stages of embryoid body formation would allow us to identify co-expressed coding and non-coding transcripts within emerging lineages. As such, we analyzed the RNAome of primed pluripotent stem cells and that of individual cells obtained from dissociated embryoid bodies at days 4, 8 and 12 of culture (**Fig. 3a**). We found that the fraction of mRNA with respect to all other analyzed transcripts was higher in pluripotent compared to differentiated cells (65% vs 50-54%) (**Fig. 3a**). Consistent with previous studies^27^, the number of coding genes expressed by pluripotent stem cells was also higher compared to differentiated progenitors (**see Supplementary Fig. 10a**). This was also the case for several non-coding RNA types, such as lncRNA, miRNA and scaRNA (**see Supplementary Fig. 10b**). Specifically, we observed that the levels of certain snoRNAs (such as *Snord17, Snora23, Snord87*), scaRNAs (such as *Scarna13* and *Scarna6*), lncRNAs (*Platr3, Lncenc1, Snhg9, Gm31659, etc.*) and miRNAs (*Mir92-2, Mir302b, Mir19b-2*) go down after cells exit pluripotency (**Fig. 3b**). In contrast, we also identified that the levels of several lncRNAs (*Tug1, Meg3, Lockd*) and miRNAs (*Mir298, Mir351, Mir370*) increase with differentiation (**Fig. 3b**).

**Figure 3.**
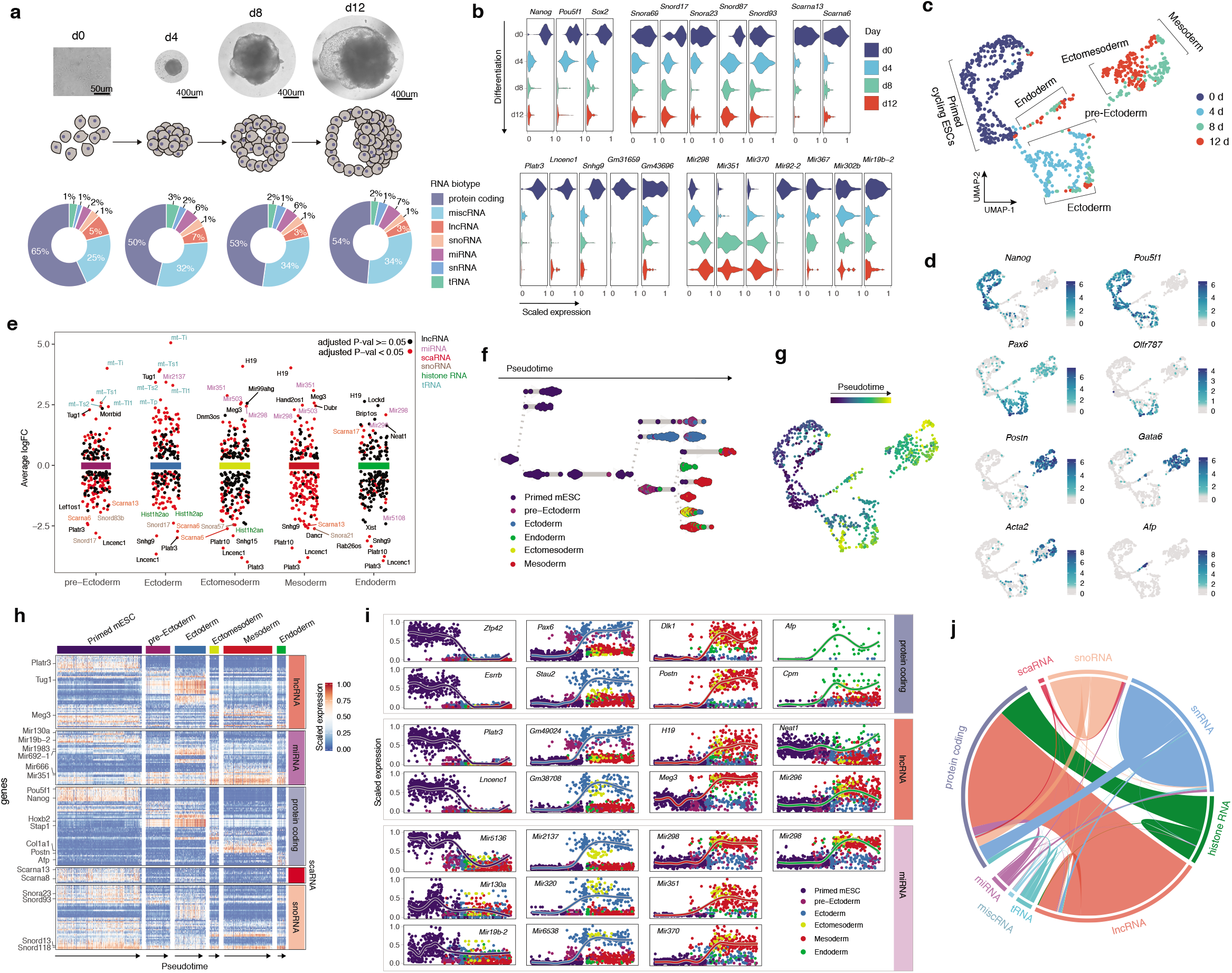
Coding and non-coding signature of differentiated single mESCs. **a. Microscope images and corresponding schematic representations of EB formation at four sampled time points.** Pie charts represent distribution of mapped reads across RNA types. Genes were assigned to a specific biotype based on GENCODE M23 annotation for the reference chromosomes. tRNA was quantified by mapping the reads, non-mapping to any other RNA type, to the high-confidence gene set obtained from GtRNAdb. **b. Exemplary coding and non-coding genes that are up- or downregulated during EB formation**. Subpanels are grouped according to RNA type. **c. UMAP plot of collected cells colored by timepoint.** Cells were clustered using k-nearest neighbor algorithm and cell lineages were annotated based on the expression of marker genes within the identified clusters. **d. UMAP plots showing the expression of lineage markers.** Color scale adjacent to each plot denotes the expression of a given gene as log1p(counts). **e. Non-coding genes differentially expressed between annotated lineages and primed mESCs.** Primed mESCs are marked by the expression of *Nanog, Pou5f1* and *Esrrb.* **f. Lineage tree of EB differentiation.** Each dot represents a cell colored according to the assigned lineage. Cells are arranged according to the computed pseudotime. **g. UMAP plot of collected cells colored by pseudotime.** **h. Heatmap showing the variability in coding and non-coding gene expression across identified clusters.** **i. Temporal and lineage-specific expression of selected protein-coding, lncRNA and miRNA genes.** Each column from left to right shows genes specific to: pluripotency state, ectoderm, mesoderm or endoderm lineages. **j. Co-expression of coding and non-coding genes in differentiating mESCs.** Gene co-expression was evaluated through pairwise correlation analysis across all cells collected at four stages of EB formation. Genes with spearman rho > 0.5 were considered co-expressed.

Louvain clustering of all collected cells revealed the presence of six molecularly distinct populations which we assigned to: primed mESC, pre-Ectoderm, Ectoderm, Endoderm, Ectomesoderm and Mesoderm (**Fig. 3c**), based on the expression of known lineage-specific marker genes (e.g. *Nanog* and *Pou5f1* for pluripotent cells, *Pax6* and *Olfr787* for ectoderm, *Afp* and *Shh* for endoderm, *Acta2* and *Col3a1* for mesoderm^28^) (**Fig. 3d, see Supplementary Fig. 11**). The analysis of genes differentially expressed between primed mESCs and each of the identified clusters showed that in addition to well-characterized lineage-specific mRNAs (**see Supplementary Fig. 12a**)^29,30^ and lncRNAs (*Tug1* in ectodermal and *Meg3* mesodermal lineages respectively)^6^, other ncRNA genes such as miRNAs, scaRNAs, snoRNAs, tRNAs and histone RNAs are either specifically expressed or downregulated within a certain lineage (**Fig. 3e**).

We next used PAGA^31^ to infer a developmental trajectory and compute pseudotime coordinates for each cell in our dataset (see Methods) (**Fig. 3f-g**). Aligning cells in pseudotime within each lineage further confirmed the existence of expression gradient within different RNA types (**Fig. 3h**). Furthermore, we found that the majority of identified variable non-coding transcripts were germ-layer specific. Examples of such transcripts include *Mir2137, Mir320, Gm49024* and *Gm38708* in ectoderm, *Mir351, Mir370* and *Meg3* in mesoderm as well as *Neat1* in endoderm. *Mir296* and *Mir298* were expressed in both mesoderm and endoderm but were absent in ectoderm (**Fig. 3i, see Supplementary Fig. 13**).

Finally, to understand the relationship between mRNA and non-coding RNA genes we performed a pairwise correlation analysis of gene expression across all sampled cells. We found that the expression of ~ 50% of identified histone-coding genes correlated with the expression of other protein-coding genes (Spearman rho > 0.5) (**Fig. 3j**). In addition, we found that multiple ncRNAs from all assayed RNA types (e.g. miRNA, snoRNA, snRNA, etc.) are positively correlated with the expression of protein-coding genes. Most of these ncRNAs represent putative uncharacterized regulators of lineage commitment.

Altogether, Smart-seq-total enables an unbiased exploration of a broad spectrum of coding and non-coding RNA transcripts in individual cells. Further improvements to Smart-seq-total can include incorporation of unique molecular identifiers, enhancement of ncRNA capture through sequence optimization or enzymatic 5’prime cap addition^32^ as well as depletion of a wider range of overrepresented RNAs. We anticipate Smart-seq-total to facilitate the identification of non-coding regulatory patterns and its functional role in regulating cellular functions and shaping cellular identity. This also means shifting the current protein-centered view of gene regulation towards comprehensive maps featuring both, protein and RNA regulators.

## Acknowledgements

We thank Giana Cirolia from CZ Biohub for her thoughtful advice regarding DASH protocol as well as for providing rRNA guides. We thank Jennifer Okamoto and Rene Sit for the assistance in sequencing of the libraries. We thank Professor Joanna Wysocka and her team members for guidance in embryoid body differentiation procedure. We thank Gita Mahmoudabadi for her constructive comments on the manuscript. This study was supported by Chan Zuckerberg Biohub. A.I. was supported by the Swiss National Foundation PostDoc Mobility Fellowship.

## Author Contributions

A.I. designed and performed the experiments and data analysis. A.I. and N.N. developed an optimized pipeline for library preparation, rRNA depletion and sequencing. A.I. and S.Q. interpreted the data and wrote the manuscript.

## METHODS

### Cell culture

HEK293T cells were cultured in complete DMEM high glucose medium (Gibco, ThermoFisher 11965092) supplemented with 5% Fetal Bovine Serum (ThermoFisher 16000044), 1mM Sodium Pyruvate (ThermoFisher 11360070) and 100 μg/mL Penicillin/Streptomycin (ThermoFisher 15070063). Human primary dermal fibroblasts were obtained from ATCC (ATCC® PCS-201-012^™^). Cells were cultured and passaged four times in Fibroblast Basal Medium (ATCC® PCS-201-030^™^) supplemented with 5ng/mL rh FGF β, 7.5mM L-glutamine, 50 ug/mL Ascorbic acid, 5ug/mL rh Insulin, and 1% Fetal Bovine Serum (Fibroblast Growth kit-low serum, ATCC^®^ PCS-201-041^™^). MCF7 cells (ATCC^®^ HTB22^™^) were cultured in complete DMEM high glucose medium (Gibco, ThermoFisher 11965092) supplemented with 10% Fetal Bovine Serum (ThermoFisher 16000044), 1mM Sodium Pyruvate (ThermoFisher 11360070) and 100 μg/mL Penicillin/Streptomycin (ThermoFisher 15070063). Cells were collected 2-4h after passaging, dissociated using 0.25% Trypsin-EDTA (ThermoFisher 25200056) and sorted in either 96-well plates containing 3uL lysis buffer or 384-well plates containing 0.3 uL of lysis buffer in each well.

mESCs were maintained and differentiated as described previously^**35,36**^. Briefly, mESCs were grown in serum-free 2i+LIF medium (complete medium: DMEM/F12 glutaMAX (Gibco, ThermoFisher 10565018), 1% N2 supplement (Gemini Bio), 2% B27 supplement (Gemini Bio), 0.05% BSA fraction V (ThermoFisher, 15260037), 1% MEM-non-essential amino acids (ThermoFisher 11140050), and 110 μM 2-mercaptoethanol (Pierce); supplemented with MEK inhibitor PD0325901 (0.8 μM), GSK3β inhibitor CHIR99021 (3.3 μM) and 10ng/mL mouse LIF (Gibco, PMC9484)) in tissue culture (TC) dishes pretreated with 7.5 μg/ml polyL-ornithine (Sigma) and 5 μg/ml laminin (BD). To induce spontaneous embryoid body formation cells were washed with PBS, dissociated with StemPro Accutase (Gibco, ThermoFisher A1110501), transferred to serum-rich medium (complete medium: DMEM/F12 glutaMAX (Gibco), 1% N2 supplement (Gemini Bio), 2% B27 supplement (Gemini Bio), 0.05% BSA fraction V, 1% MEM-non-essential amino acids, and 110 μM 2-mercaptoethanol; supplemented with 10% FBS (ThermoFisher 10439001)) and diluted to 10^6 cells/mL. Each 10 uL of cell suspension were plated as a hanging drop in 10 cm^2^ TC dishes (15-20 drops per dish). 10uL of fresh serum-rich media was added to each drop on the day 4 post seeding. Primed mESCs were collected 6h after seeding. Embryoid bodies were collected and dissociated at days 4, 8 and 12 of culture.

### Cell sort

Lysis plates were prepared by dispensing 0.3μL lysis buffer (4 U Recombinant RNase Inhibitor (RRI) (Takara Bio, 2313B), 0.12% Triton^™^ X-100 (Sigma, 93443-100ML), 1μM Smart-seq-total oligo-dT primer (5’-Biotin-CATAGTCTCGTGGGCTCGGAGATGTGTATAAGAGACAGT30VN-3’;IDT) (see **Supplementary Table 3** for a full list of oligos used in the present study) into 384-well hard-shell PCR plates (Bio-Rad HSP3901) using Mantis liquid handler (Formulatrix). 96-well lysis plates were prepared with 3 μl lysis buffer. All plates were sealed with AlumaSeal CS Films (Sigma-Aldrich Z722634), spun down and snap-frozen on dry ice.

Cells were stained with calcein-AM and ethidium homodimer-1 (LIVE/DEAD^®^ Viability/Cytotoxicity Kit, ThermoFisher L3224) and individual live cells were sorted in 384 well lysis plates using SONY sorter (SH800S) with 100um nozzle chip. Plates were spun down and stored at −80 degrees immediately after sorting.

### Generation of Smart-seq-total libraries

To facilitate cell lysis and denaturation of the RNA, plates were incubated at 72 degrees for 3 min, and immediately placed on ice afterwards. Next, 0.2 uL of polyA tailing mix, containing 1.25U E.coli PolyA (NEB M0276S), 1.25X PolyA buffer (NEB), 1.25 mM ATPs (NEB) and 4U of RRI (Takara); were added to each samples. PolyA tailing was carried out for 15 minutes at 37 °C followed by 72 °C for 4 minutes. After polyA tailing plates were immediately placed on ice for 2-5 minutes. 1uL of reverse transcription mix, containing 15U SuperScript II (ThermoFisher), 4U RRI (Takara), 1.5X First-Strand Buffer, 1.5 μM TSO (Exiqon, 5’-biotin-UCGUCGGCAGCGUCAGUUGUAUCAACUCAGACAUrGrG+G-3’), 7.5 mM DTT, 1.5 M Betaine (Sigma, B0300-5VL), 10 mM MgCl2 (Sigma, M1028-10X1ML) and 1.5 mM dNTPs (ThermoFisher, 18427013); was added to each well. Reverse transcription was carried out at 42 °C for 90 min, and terminated by heating at 85 °C for 5 min. Subsequently, 0.3 uL of TSO digestion buffer containing 1U Uracil-DNA glycosylase (UDG, NEB M0280S) were added to each well. Plates were incubated for 30 minutes at 37 °C. PCR preamplification was performed directly after TSO digestion by adding 3.2 μL of PCR mix to each well, bringing the reaction concentration to 1x KAPA HiFi MIX (Roche), 0.5 μM Forward PCR primer (5’-TCGTCGGCAGCGTCAGTTGTATCAACT-3’; IDT), 0.5 μM Reverse PCR primer (5’-GTCTCGTGGGCTCGGAGATGTG-3’; IDT). PCR was cycled as follows: 1) 95 °C for 3 min, 2) 21 cycles of 98 °C for 20 s, 67 °C for 15 s and 72 °C for 6 min, and 3) 72 °C for 5 min. The amplified product was cleaned up using 1X ratio of AMPure beads on Bravo liquid handler platform (Agilent). Concentrations of purified product were measured with a dye-fluorescence assay (Quant-iT PicoGreen dsDNA High Sensitivity kit; Thermo Fisher, Q33120) on a SpectraMax i3x microplate reader (Molecular Devices). Samples were then diluted to 0.2 ng/uL. To generate sequencing libraries, 1.5uL of diluted samples was amplified in a final volume of 5uL using 2X KAPA mix and 0.4 μl of 5 μM i5 indexing primer, 0.4 μl of 5 μM i7 indexing primer. PCR amplification was carried out using the following program: 1) 95 °C for 3 min, 2) 8 cycles of 98 °C for 20 s, 65 °C for 15 s and 72 °C for 1 min, and 3) 72 °C for 5 min.

### Library pooling, ribosomal sequence digestion and sequencing

After library preparation, wells of each library plate were pooled using a Mosquito liquid handler (TTP Labtech). Pooling was followed by a purification with 0.8x AMPure beads (Fisher, A63881). Ribosomal reads were digested using DASH as described in ^33^. Briefly, 135 guides designed to target 45S rRNA sequence (Supplementary Table 1) were combined with tracer RNA and assembled with Cas9 protein in 2:1 ratio. The assembled complexes were incubated with the sequencing library in 1X Cas9 buffer (Supplementary Table 1) for 1h at 37°C. Following rRNA sequence digestion Cas9 was inactivated through incubation with proteinase K for 15min at 50°C. Library was then purified twice, first using 1.2X and then 0.8x AMPure beads:DNA ratio. Library quality was assessed using capillary electrophoresis on a Fragment Analyzer (AATI), and libraries were quantified by qPCR (Kapa Biosystems, KK4923) on a CFX96 Touch Real-Time PCR Detection System (Biorad). Plate pools were normalized to 2 nM and equal volumes from 8 plates were mixed together to make the sequencing sample pool. A PhiX control library was spiked in at 10% before sequencing. Libraries were sequenced on the NovaSeq 6000 Sequencing System (Illumina) using 1 × 75 or 1×100-bp single-end reads (using custom Read 1 sequencing primer:5’-TCGGCAGCGTCAGTTGTATCAACTCAGACATGGG-3’) and 2 × 12-bp index reads.

### Data processing

Sequences from the NovaSeq were de-multiplexed using bcl2fastq version 2.19.0.316. Reads were trimmed from polyA tails using cutadapt v 1.18 with the following parameters: -m 18 -j 4 -a AAAAAAAAAA -a TTTTTTTTTT. Reads were then aligned to the human (GRCh38) or mouse (GRCm38) genomes using STAR_v2.7.0d^37^ with the following parameters --outFilterMismatchNoverLmax 0.05 --outFilterMatchNmin 18 --outFilterMatchNminOverLread 0 --outFilterScoreMinOverLread 0 --outMultimapperOrder Random. Reads mapping to multiple locations were assigned either to a location with the best mapping score or, in the case of equal multimapping score – to the genomic location randomly chosen as “primary”.

Transcripts were counted using *featureCounts v 1.6.1* ^38^ with the following parameters -M –primary -s 1. GENCODE v32 and GENCODE M23^39^ annotations were used for human and mouse reads respectively. tRNA was quantified using high-confidence gene set obtained from GtRNA^40^. To account for multimappers “primary” alignment reported by STAR was counted. For miRNA and tRNA all reads mapping to arms or the stem loop were summed to quantify the expression at the gene level.

### Comparison of Smart-seq2 and Smart-seq-total

HEK293T cells were sorted in 96-well plates containing 3uL of lysis buffer (as described above). The reaction volumes for Smart-seq-total were scaled 10 times compared to 384-plate format, i.e. RNA from each cell was polyadenylated in 5uL, reverse transcribed in 15 uL and cDNA was pre-amplified in 15uL total volume. We retrieved Smart-seq2 data from (Picelli et al., 2013; GSE49321). Smart-seq2 and Smart-seq-total reads were mapped using STAR and counted using featureCounts as described above. Comparisons between protocols in Fig 1b were generated on depth-normalized libraries, using 2.5 million randomly selected reads per adaptor-trimmed library (or all reads for libraries that had less than 2.5 million reads) to compute expression levels (cpm).

### Unsupervised clustering and dimensionality reduction analysis of human cell types

Standard procedures for filtering, variable gene selection, dimensionality reduction and clustering were performed using the Seurat package version 3.1.4^41^. Cells with fewer than 2000 detected genes and those with more than 2 Mio reads were excluded from the analysis. Counts were log-normalized for each cell using the natural logarithm of 1 + counts per million. Variable genes were selected based on overdispersion analysis and projected onto a low-dimensional subspace using principal component (PC) analysis. The number of PCs was selected on the basis of inspection of the plot of variance explained. Cells were visualized using a 2-dimensional t-distributed Stochastic Neighbor Embedding of the PC-projected data. Dimensionality reduction parameters for tSNE (resolution and number of PCs) were adjusted on a per-cell type and per-biotype basis and can be viewed in the Rmd files available on GitHub. Cells were assigned a cell cycle score using Seurat’s CellCycleScoring() function using cell cycle markers described in ^42^.

### Clustering of coding and non-coding genes

Clusters of coding and non-coding genes shown in Fig. 2b were computed and visualized using DEGreport R package^43^. Top 250 marker genes for each cell cycle phase and all non-coding genes with average expression ln(cpm+1)> 0.05 in at least one phase were used for this analysis. Gene expression values were normalized using variance stabilizing transformation^44^ before clustering. Further details of the analysis can be viewed in the Rmd files available on GitHub.

### Pre-processing and clustering of mESCs

Standard procedures for filtering, variable gene selection, dimensionality reduction and clustering were performed using the Seurat package version 3.1.4^41^. Cells with fewer than 1000 detected genes and those with more than 2 Mio reads were excluded from the analysis. Counts were log-normalized for each cell using log1p(counts) and 1e4 scale factor. Variable genes were projected onto a low-dimensional subspace using PC analysis. The number of PCs was selected on the basis of inspection of the variance explained plot. A shared-nearest-neighbor graph was constructed on the basis of the Euclidean distance in the lowdimensional subspace spanned by the top PCs. Cells were visualized using Uniform manifold Approximation and Projection (UMAP) algorythm^45^ of the PC-projected data. Clusters were annotated based on the expression of known marker genes corresponding to one of the three germ layers. Cells were assigned a cell cycle score using Seurat’s CellCycleScoring() function and cell cycle markers described in^42^.

### Developmental trajectory inference of EB differentiation

Developmental trajectory of mESC differentiation was inferred using PAGA through dynoverse wrapper^46^. Pseudotime coordinates computed from the trajectory were appended to Seurat object and further used to generate Figures 1f-i.

### Correlation between coding and non-coding RNA levels

Spearman coefficients were computed for all pair-wise correlations of expressed genes (average ln(cpm+1)>2 across all cells). The resulting matrix was subset to only mRNA:non-coding RNA correlations. Pairs with Spearman rho > 0.5 were used to generate a chord diagram shown in Figure 3h and pairs with Spearman rho < −0.5 were used to generate a chord diagram shown in Supplementary Figure 12b.

### Code availability

All code used for analysis is available on GitHub (https://github.com/aisakova/smart-seq-total/).

### Data availability

The datasets generated and analyzed in the study are available in the NCBI Gene Expression Omnibus (GEO) under the entry GEO: GSE151334.

