## Supplemental figures and tables for "Single cell profiling of total RNA using Smart-seq-total"

#### Overview:

##### Supplementary figures:

Supplementary figure 1. Smart-seq-total performance.

Supplementary figure 2. Number of coding and non-coding genes detected in primary fibroblasts, HEK293T and MCF7 cells.

Supplementary figure 3. Most abundant transcripts detected in primary fibroblasts, HEK293T and MCF7 cells.

Supplementary figure 4. Cell type-specific marker genes grouped by RNA type.

Supplementary figure 5. Transcripts dynamically changing over the cell cycle identified in primary fibroblasts.

Supplementary figure 6. Transcripts dynamically changing over the cell cycle identified in HEK293T cells.

Supplementary figure 7. Transcripts dynamically changing over the cell cycle identified in MCF7 cells.

Supplementary figure 8. Clusters of coding and non-coding genes dynamically expressed throughout the cell cycle.

Supplementary figure 9. Cell cycle bias in cell clustering.

Supplementary figure 10. Number of coding and non-coding genes detected in mESCs at each stage of EB formation.

Supplementary figure 11. UMAP plots of lineages and key marker genes.

Supplementary figure 12. Genes differentially expressed between lineages and correlation analysis.

Supplementary figure 13. miRNAs expressed in the profiled mESCs and differentiated progenitors.

##### Supplementary tables:

Supplementary table 1. CRISPR guides for rRNA depletion.

Supplementary table 2. Cell cycle-specific gene clusters.

Supplementary table 3. Smart-seq-total primers used in the present study.

**Supplementary figure captions:**

**Supplementary figure 1. Smart-seq-total performance.**

**a.** Sequencing scheme of a scRNA-seq library prepared with Smart-seq-total.

**b.** Bioanalyzer traces showing fragment sizes of amplified cDNA prepared from single cells using Smart-seq-total with and without TSO removal step.

**c.** Mean number of genes per biotype detected by Smart-seq2 and Smart-seq-total in single HEK293T cells. Same as **Fig. 1b** but for tRNA, miscRNA, TEC, intronic and antisense transcripts.

**d.** Read coverage of multiple gene types shown for Smart-seq2 and Smart-seq-total data. Computed as a sum of n=4 cells.

**Supplementary figure 2. Number of coding and non-coding genes detected in primary fibroblasts, HEK293T and MCF7 cells.**

**a.** Number of counts and number of genes per cell grouped by RNA type. Computed based on 612 profiled cells.

**b.** Number of genes, number of counts as well as the percentage of mitochondrial and histone RNA per cell computed for primary fibroblasts, HEK293T and MCF7 cells.

**c.** Number of detected genes by RNA type in the three profiled cell types.

**Supplementary figure 3. Most abundant transcripts detected in primary fibroblasts, HEK293T and MCF7 cells.**

Top 100 genes in each of the three profiled cell types supported by the largest number of reads. Red arrows indicate cell type-specific genes.

**Supplementary figure 4. Cell type-specific marker genes grouped by RNA type.**

**a.** Dot plots of marker genes identified in primary fibroblasts, HEK293T and MCF7 cells grouped by RNA type.

**b.** Relative levels of 21 tRNA (iMet-tRNA) types identified in fibroblasts, HEK293T and MCF7 cells as a fraction of total reads mapped to tRNA genes.

**Supplementary figure 5. Transcripts dynamically changing over the cell cycle identified in primary fibroblasts.**

Coding and non-coding RNAs differentially expressed across the cell cycle in primary dermal fibroblasts. Grouped by RNA type. Circular charts depict average expression of a given gene across all cells identified to be in a certain phase of the cell cycle.

**Supplementary figure 6. Transcripts dynamically changing over the cell cycle identified in HEK293T cells.**

Same as Supplementary fig. 5 but for HEK293T cells.

**Supplementary figure 7. Transcripts dynamically changing over the cell cycle identified in MCF7 cells.**

Same as Supplementary fig. 5-6 but for MCF7 cells.

**Supplementary figure 8. Clusters of coding and non-coding genes dynamically expressed throughout the cell cycle.**

Universal and cell-type specific gene clusters composed of coding and non-coding genes that dynamically change through the cell cycle. Hierarchical clustering was performed on a mixed-gene group composed of top 750 mRNAs differentially expressed through the cell-cycle and non-coding genes expressed in least one phase. For more information on cluster composition see Supplementary table 2.

**Supplementary figure 9. Cell cycle bias in cell clustering.**

**a.** t-SNE plot of the three profiled human cell types generated based on all detected genes.

**b.** t-SNE plot of the three profiled human cell types colored by cell cycle phase.

**c.** t-SNE plots colored by expression of selected cell-cycle and histone genes. Color scale adjacent to each plot denotes the expression of a given gene as ln(cpm+1).

**Supplementary figure 10. Number of coding and non-coding genes detected in mESCs at each stage of EB formation.**

**a.** Total number of detected genes and counts, as well as the percentage of mitochondrial and histone RNA, detected in each cell collected at day 0, day4, day 8 and day 12 of embryoid body formation.

**b.** Number of counts and number of genes detected in each cell at days 0, 4, 8 and 12 of embryoid body formation. Grouped by RNA type.

**Supplementary figure 11. UMAP plots of lineages and key marker genes.**

**a.** UMAP plot of collected murine cells colored by annotated cell types.

**b.** UMAP plots showing the expression of lineage markers. Color scale adjacent to each plot denotes the expression of a given gene as log1p(counts).

**a.** UMAP plot of collected murine cells colored by cell cycle phase.

**Supplementary figure 12. Genes differentially expressed between lineages and correlation analysis.**

**a.** Genes differentially expressed between annotated lineages and primed mESCs.

**b.** Anticorrelation of coding and non-coding genes. Evaluated through pairwise gene correlation analysis across all cells. Genes with spearman rho < -0.5 were considered to be anticorrelated.

**Supplementary figure 13. miRNAs expressed in the profiled mESCs and differentiated progenitors.**

UMAP plots showing the expression of miRNAs in profiled cells. Color scale adjacent to each plot denotes the expression of a given gene as log1p(counts).

Supplementary figure 1

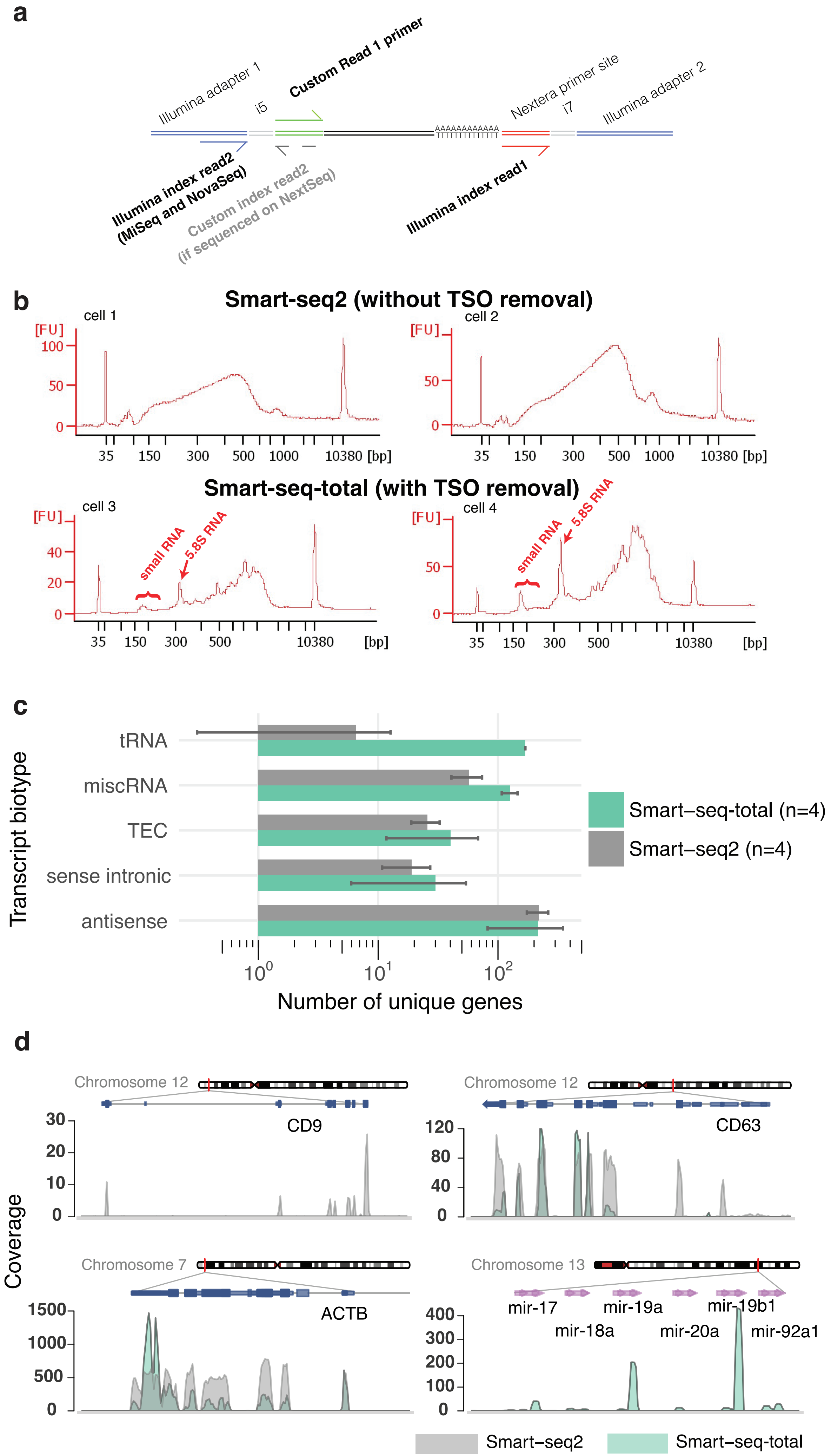

Supplementary figure 2

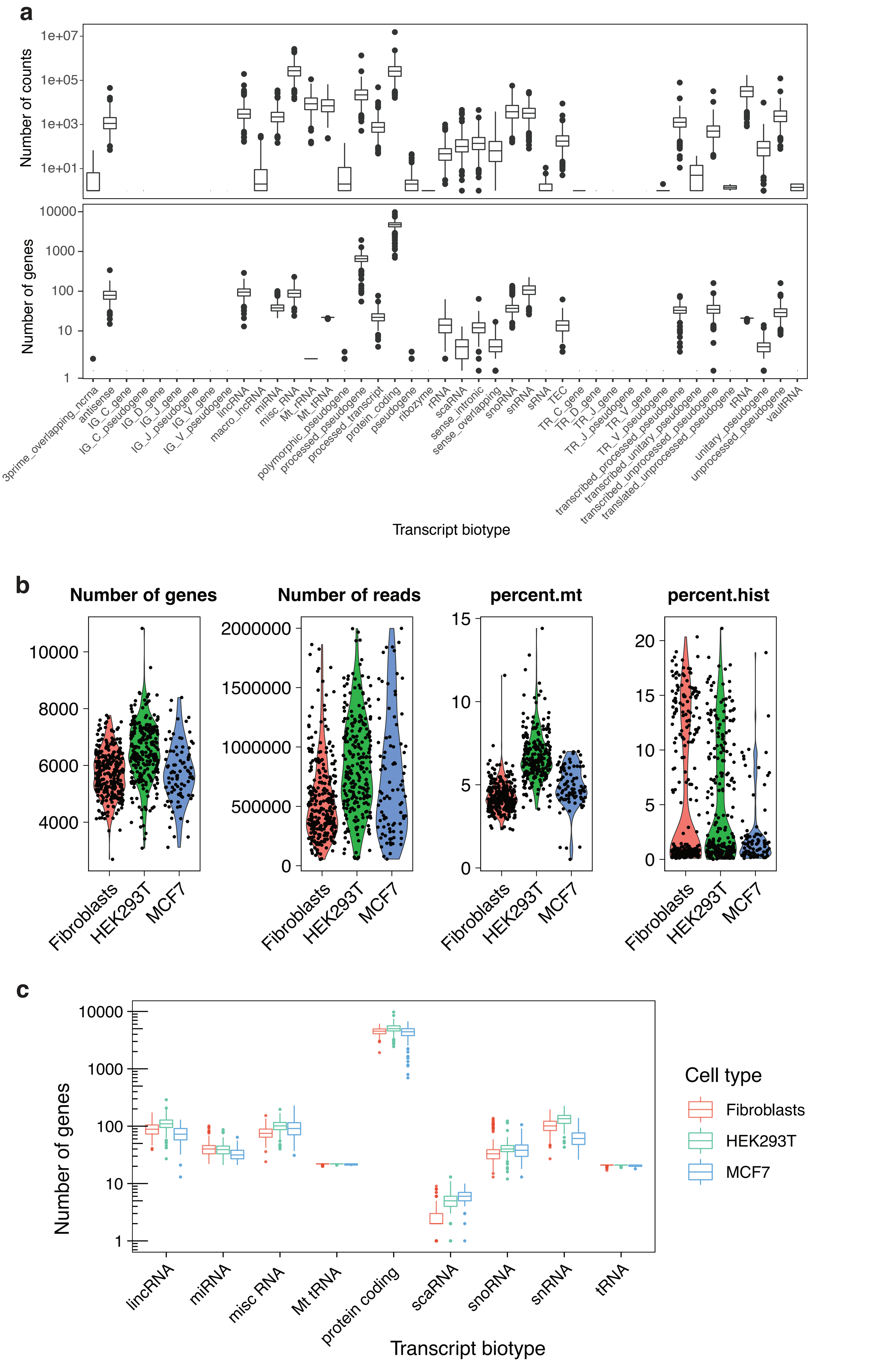

Supplementary figure 3

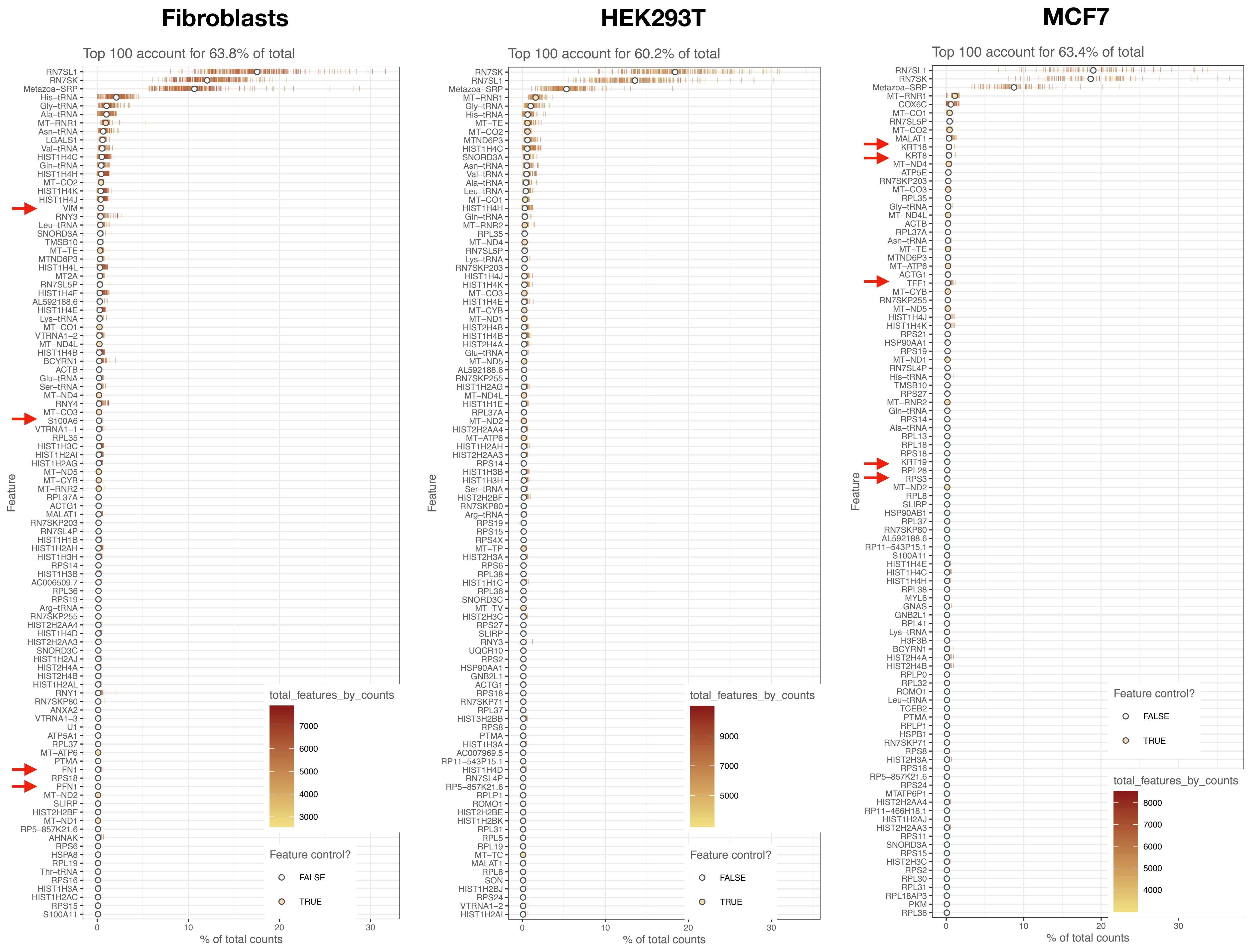

#### Supplementary figure 4

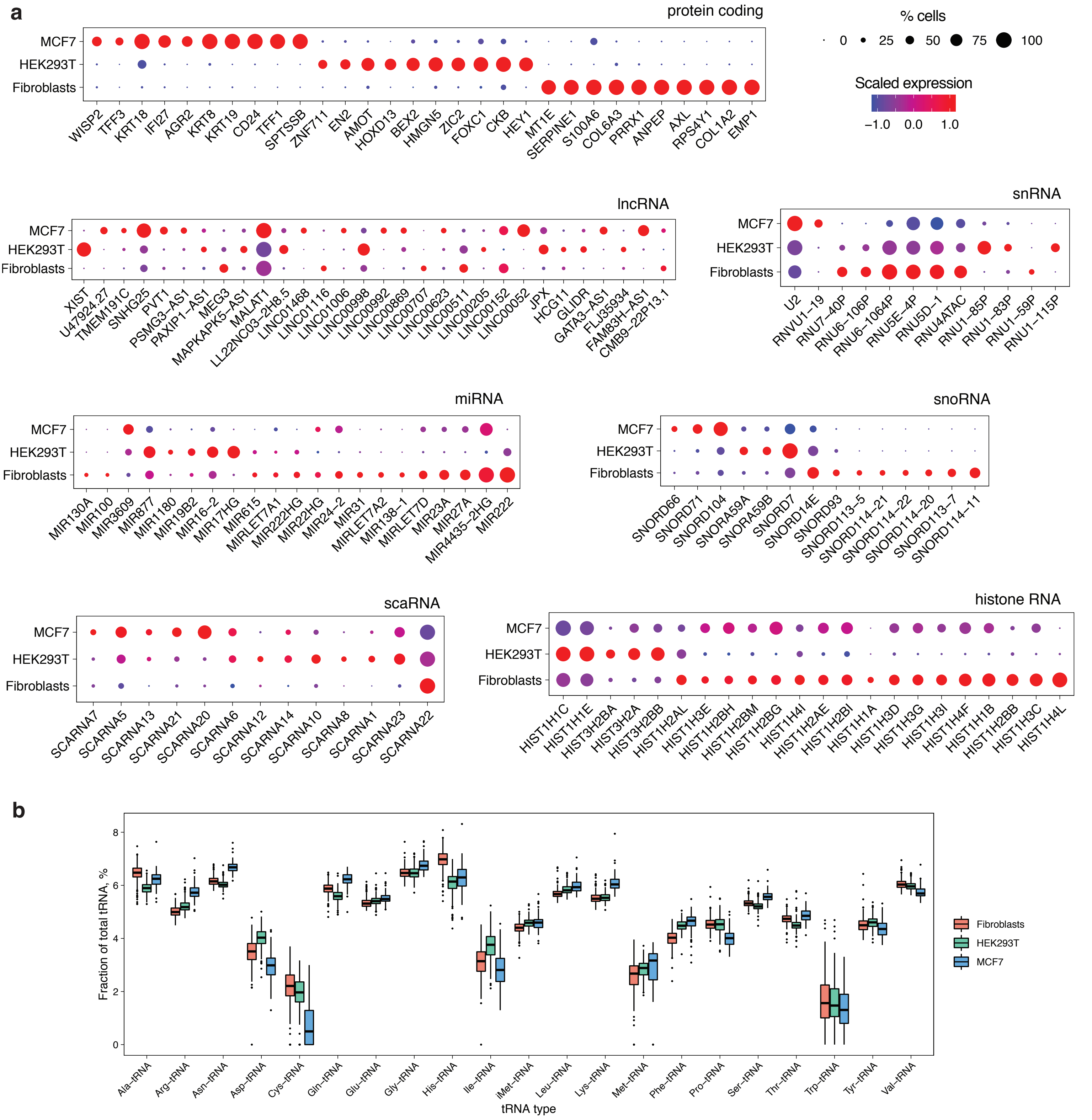

### Supplementary figure 5

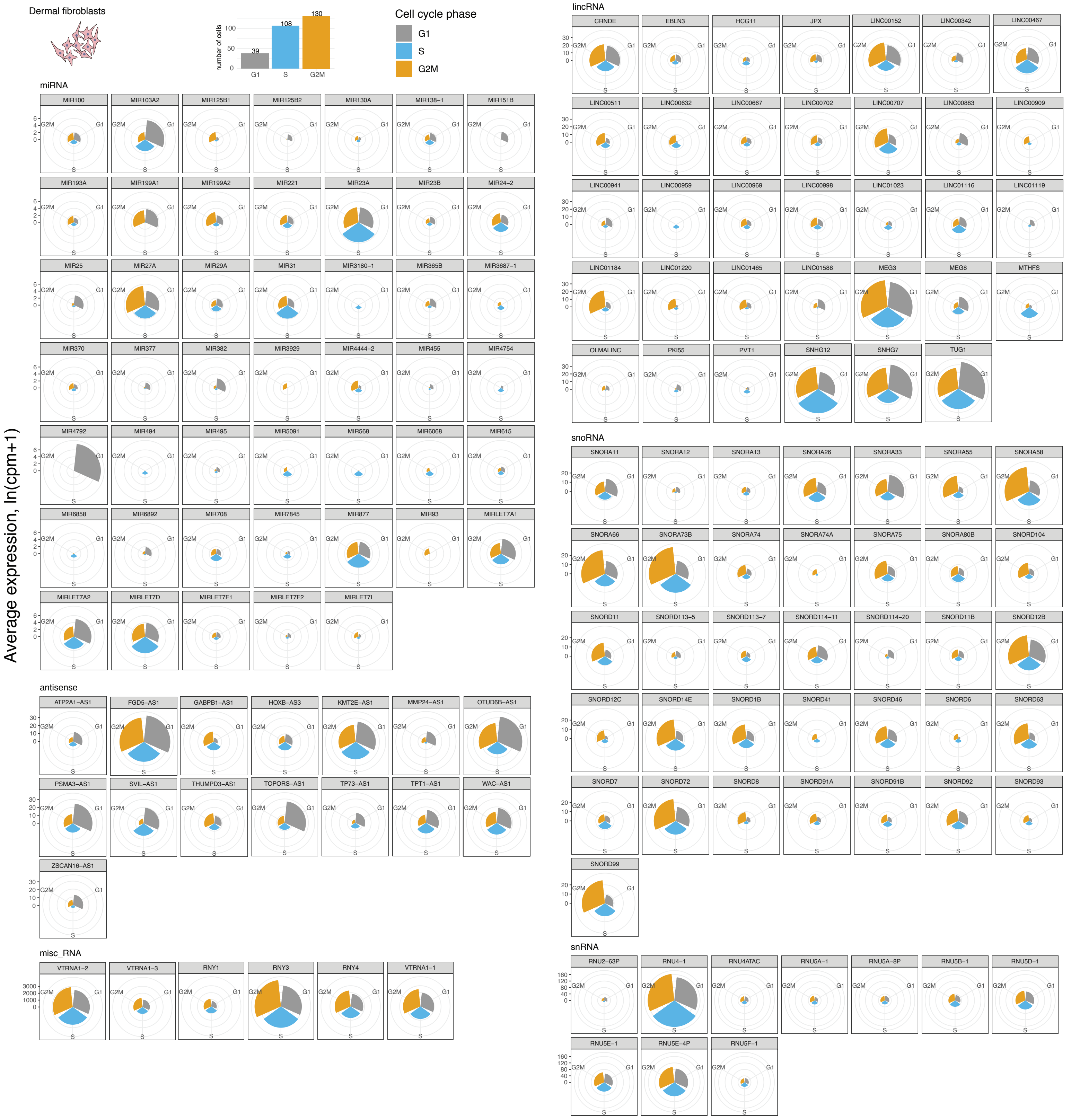

### Supplementary figure 6

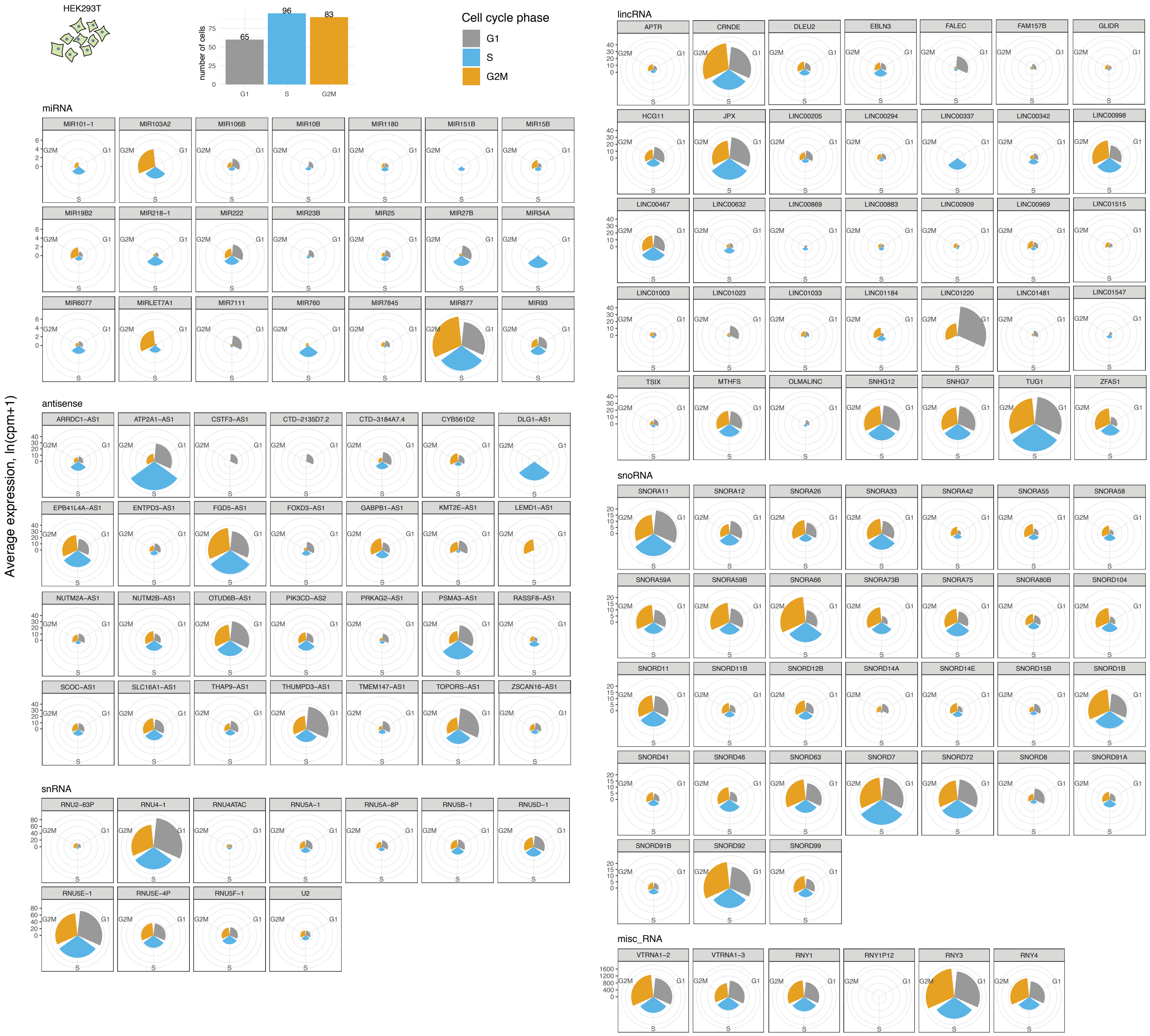

### Supplementary figure 7

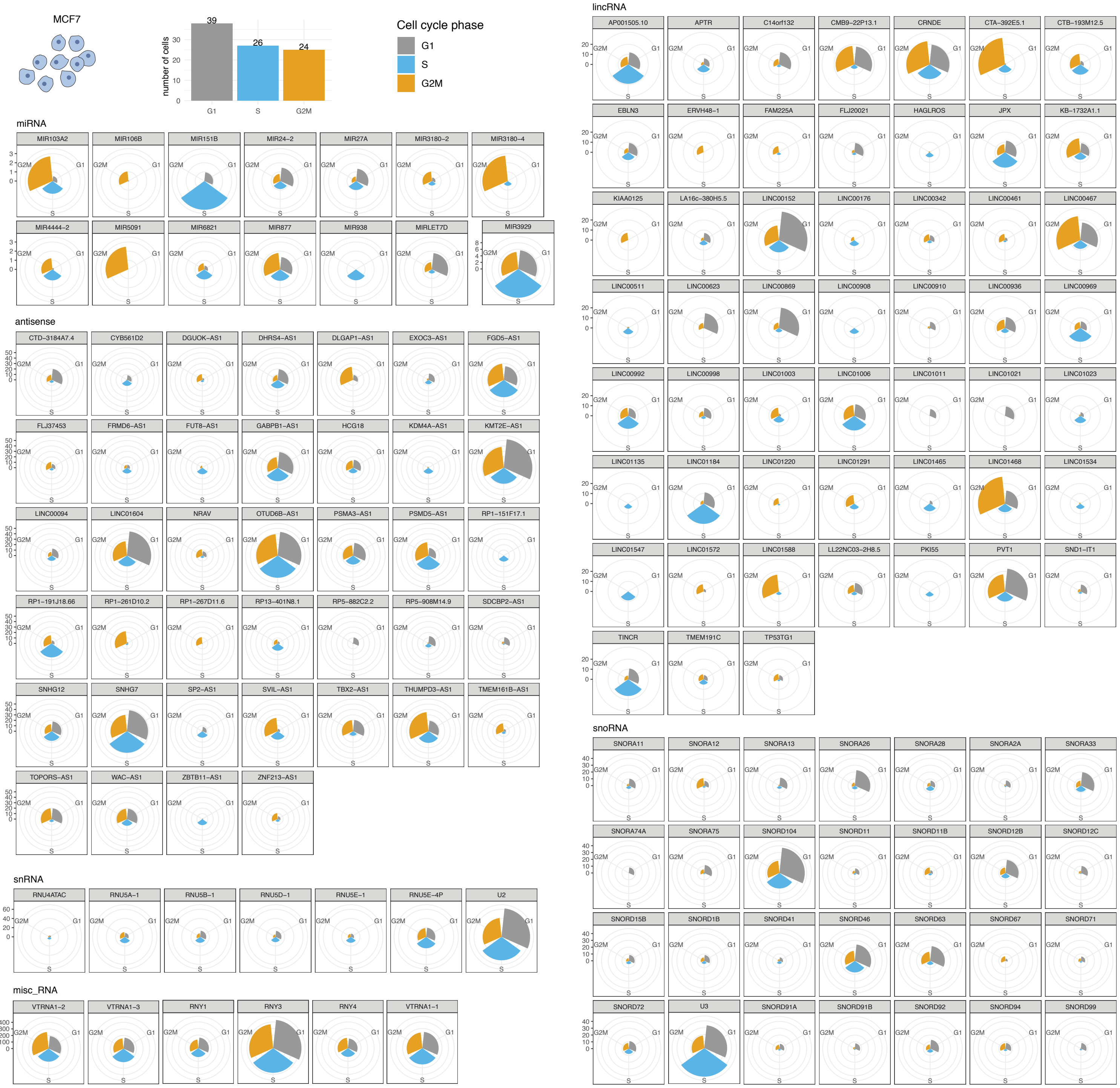

Supplementary figure 8

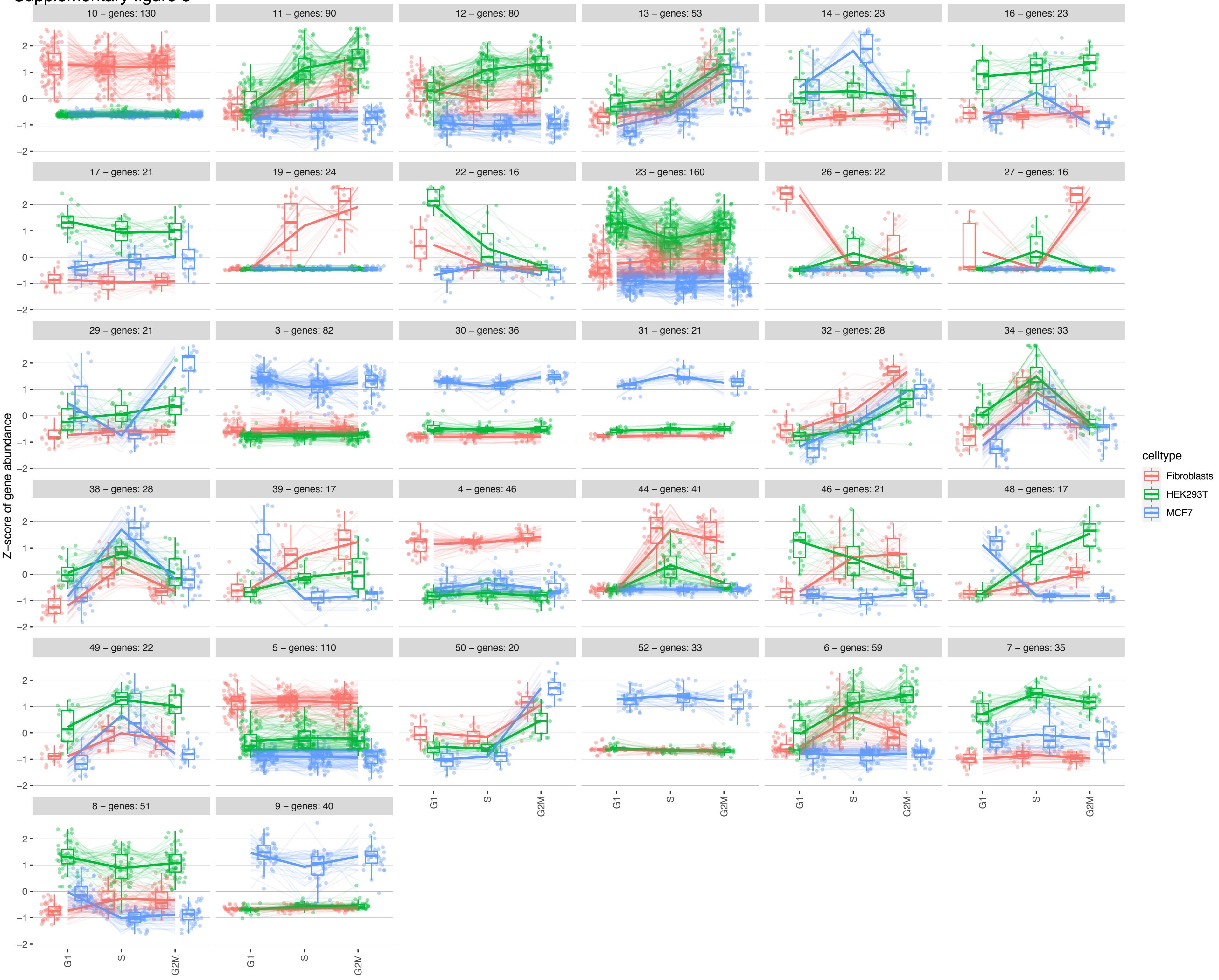

Supplementary figure 9

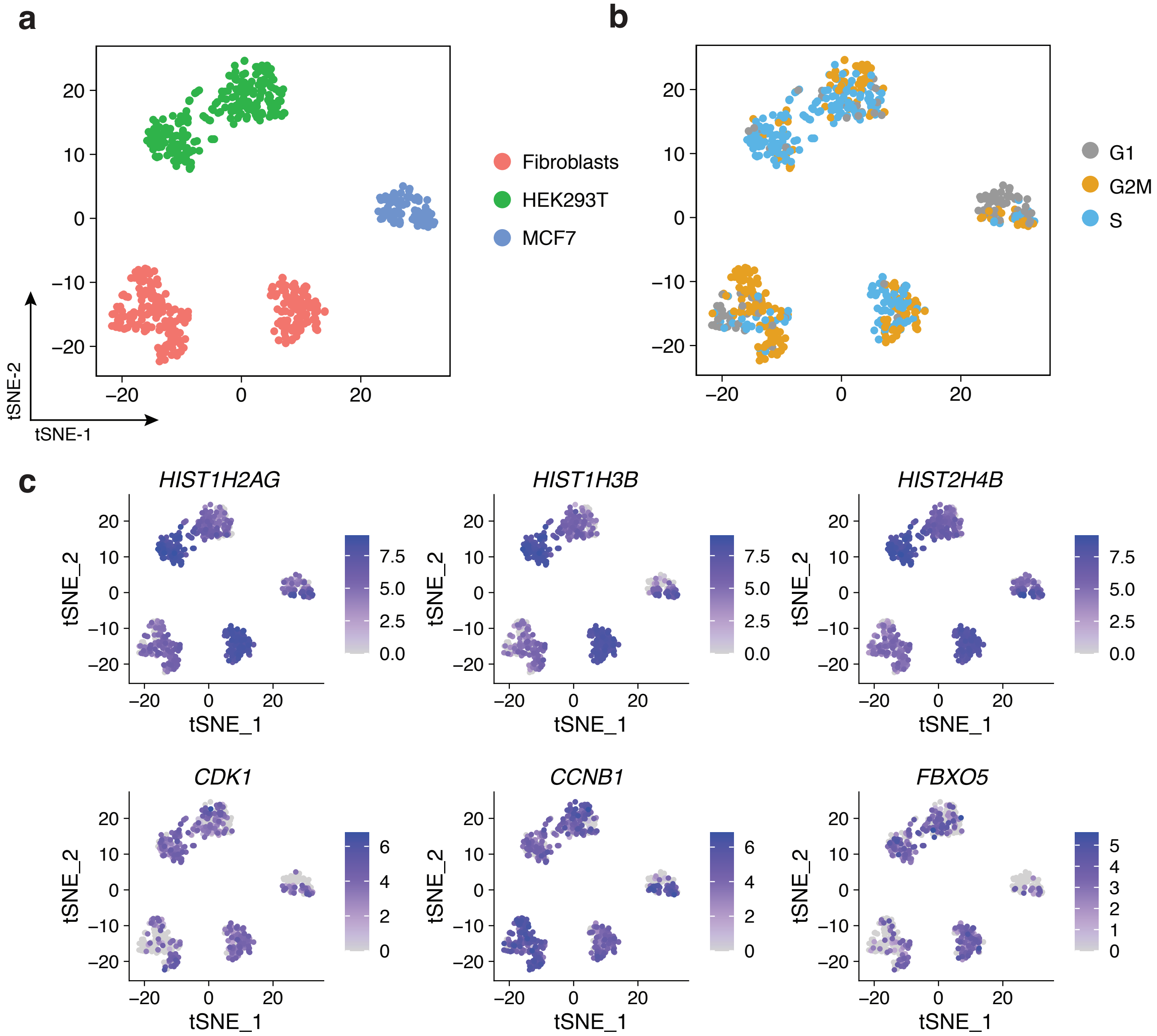

Supplementary figure 10

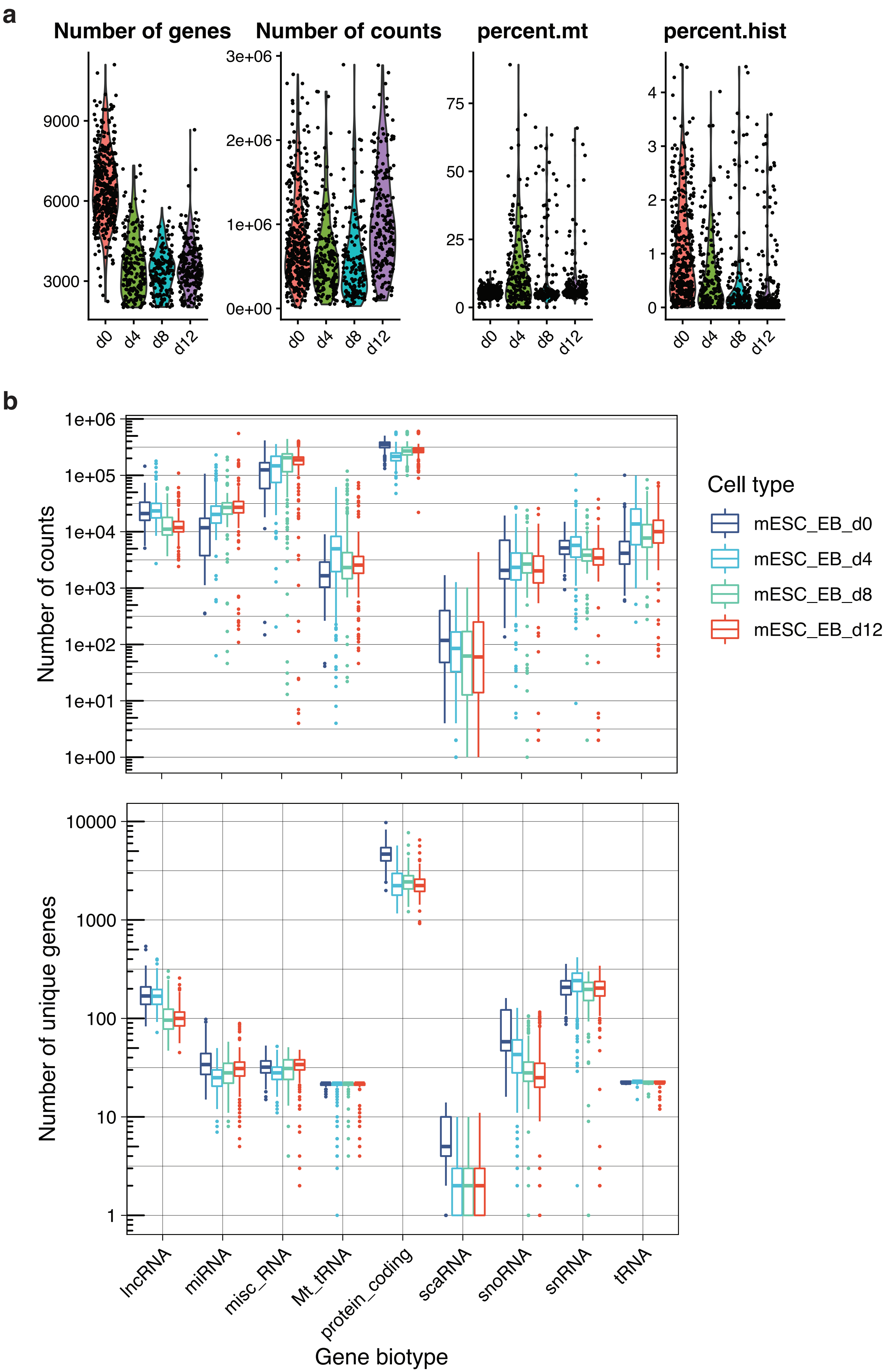

Supplementary figure 11

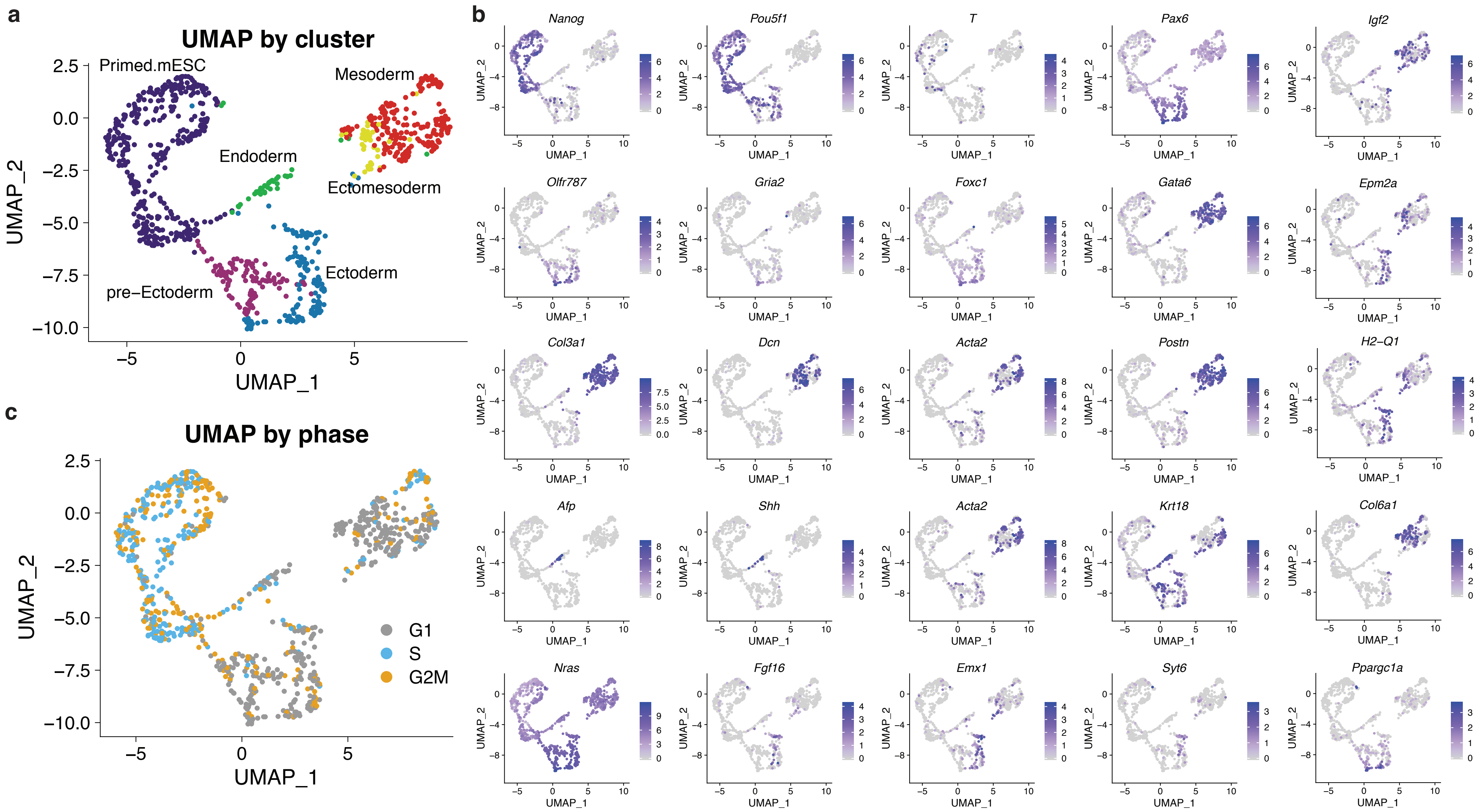

Supplementary figure 12

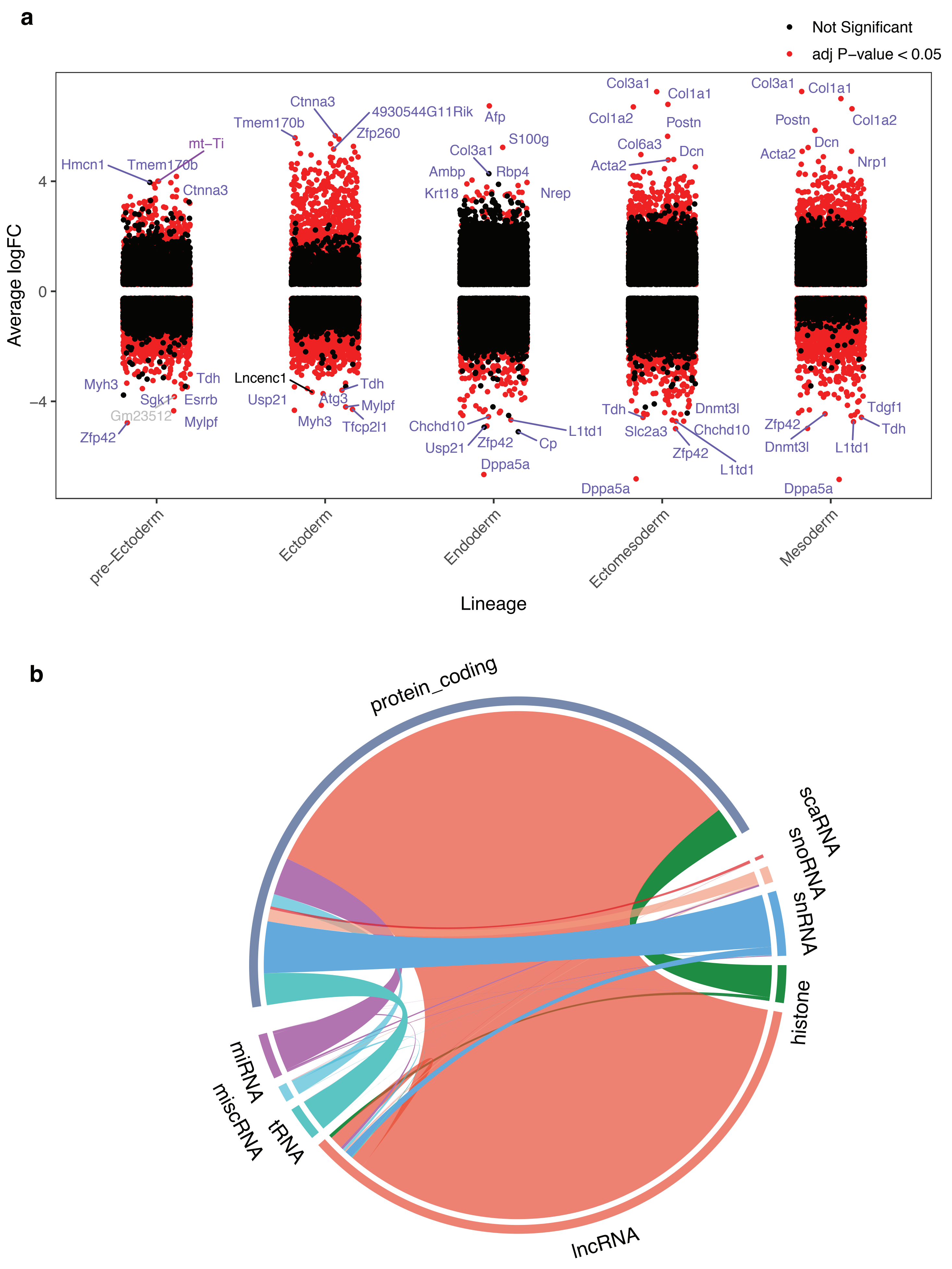

Supplementary figure 13

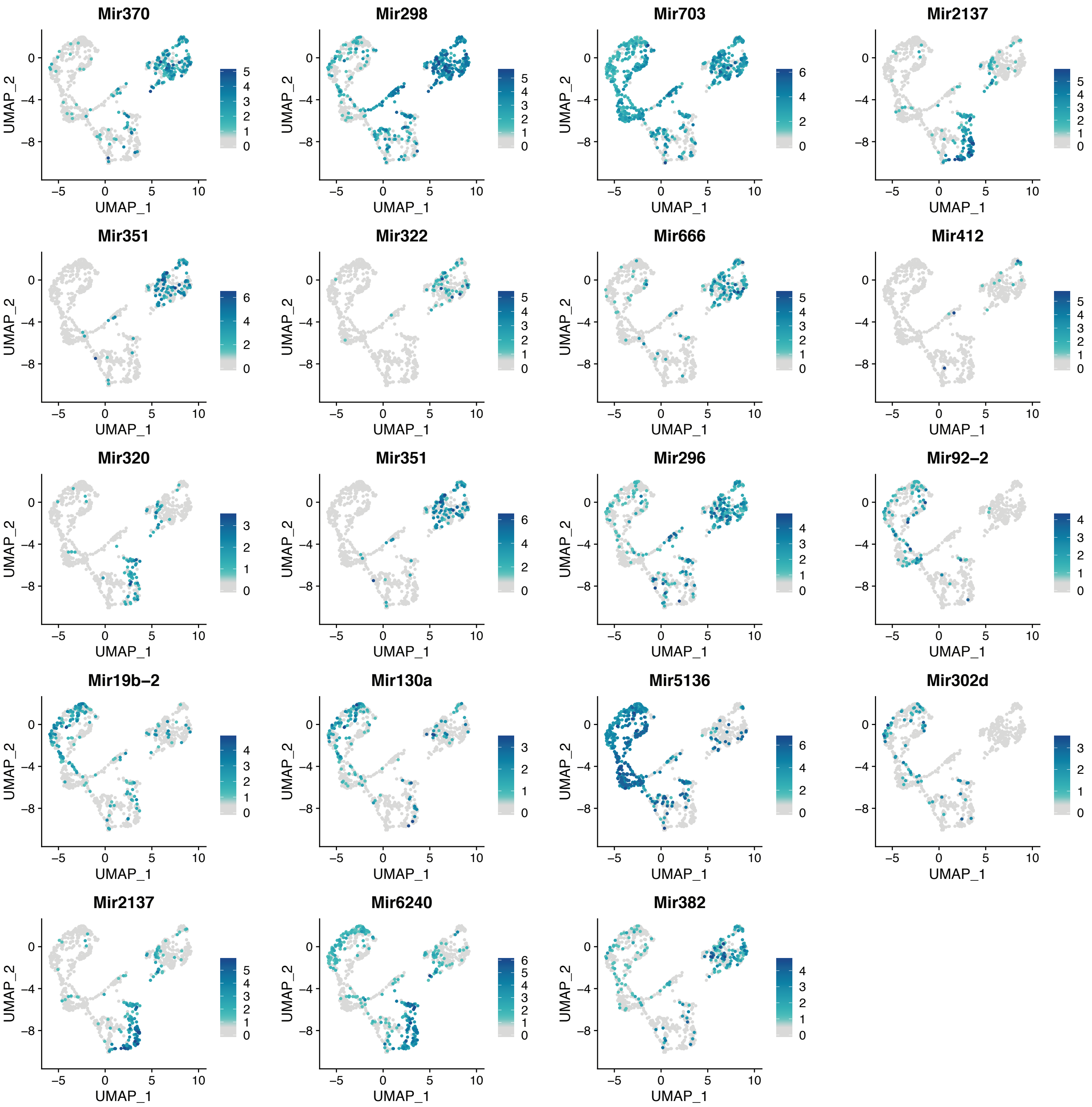

**Supplementary tables:**

Supplementary table 1. CRISPR guides for rRNA depletion.

| Guide Name | Guide Sequences (5'-3' excluding PAM sequence) |
|---|---|
| RN45s_crRNA1 | UUUAACGAGGAUCCAUUGGA |
| RN45s_crRNA2 | GUCUUGUGUGUGUCCUCGCC |
| RN45s_crRNA3 | CCGCCUCGGGGGCGUGCGUU |
| RN45s_crRNA4 | CGCGCCGGUUCGCGUCUGCU |
| RN45s_crRNA5 | AGGCAUCGGUGUGUCGGCAU |
| RN45s_crRNA6 | CGCCUGUGCGCCUGUGCGUA |
| RN45s_crRNA7 | CGCCCGGUCGCGUUCGUGCC |
| RN45s_crRNA8 | CAAAUUACCCACUCCCGACC |
| RN45s_crRNA9 | GCCGUGUGCCCGCGCGGUGU |
| RN45s_crRNA10 | GCUGGCCCGCGUCGCGGGUG |
| RN45s_crRNA11 | GGUCUGGGGGCGUGCCCUCG |
| RN45s_crRNA12 | CGGGUCUGUGUGCGCGUGUG |
| RN45s_crRNA13 | CAUGCAUGUCUAAGUACGCA |
| RN45s_crRNA14 | UGCCCUCACGUGUUUCACUU |
| RN45s_crRNA15 | GGGUUUUUGACCCGUCCCGG |
| RN45s_crRNA16 | GCUGGGGCGGUUGUCGCGUG |
| RN45s_crRNA17 | UUCGGGCCCCUCCCGGUUGG |
| RN45s_crRNA18 | UCGUAGUUGGAUCUUGGGAG |
| RN45s_crRNA19 | GAGUUCGGGGAGGGAUCACG |
| RN45s_crRNA20 | CCCUUCUCGCGCCUUCCCGU |
| RN45s_crRNA21 | GAGGAAUUCCCAGUAAGUGC |

| | |
|---|---|
| RN45s_crRNA22 | CUGACCUCGCCACCCUACCG |
| RN45s_crRNA23 | UCCUUCCAUCUCUCGCGCAA |
| RN45s_crRNA24 | CGUUAUUCCCAUGACCCGCC |
| RN45s_crRNA25 | CGGACAUUCAUGGCGAAUGG |
| RN45s_crRNA26 | GGGGUCGGUCUGGGUCCGUC |
| RN45s_crRNA27 | CGGGGACGGGACCGUUCUGU |
| RN45s_crRNA28 | UCCGAUAACGAACGAGACUC |
| RN45s_crRNA29 | CUCUCCUACUUGGAUAACUG |
| RN45s_crRNA30 | AAGGCAGGGGUGCGGCUCUC |
| RN45s_crRNA31 | GUGGCGCUUCGUCGGGGUUC |
| RN45s_crRNA32 | GCGGUUCGGGUGUGUGCUUG |
| RN45s_crRNA33 | UUAUCAGAUCAAAACCAACC |
| RN45s_crRNA34 | UUCGUCGUCCGCUCCGGGCG |
| RN45s_crRNA35 | CUUUCUCUCUGUGGGCGGGU |
| RN45s_crRNA36 | CUCUUUCUCGAUUCCGUGGG |
| RN45s_crRNA37 | AGGGAAGUCGGUCGUUCGGG |
| RN45s_crRNA38 | UCUGAGAAGCCCGUGAGAGG |
| RN45s_crRNA39 | CAUUAAUCAAGAACGAAAGU |
| RN45s_crRNA40 | CGGCUAGACGCGGGUGUCGC |
| RN45s_crRNA41 | CGCGGCAGCGUUCCCACGGC |
| RN45s_crRNA42 | CUGUGGUUUGGAGGGCGUCC |
| RN45s_crRNA43 | UUUAGUGAGGCCCACGGCCC |
| RN45s_crRNA44 | CGUGUCCCGGUGUGGCGGUG |
| RN45s_crRNA45 | AGGUGUCGGAGAGCUGUCCC |

| | |
|---|---|
| RN45s_crRNA46 | AGGUGUGGUGGGACUGCUCA |
| RN45s_crRNA47 | ACCCGAGAUUGAGCAAUAAC |
| RN45s_crRNA48 | AAAGGGAAAGAGGCUAGCAG |
| RN45s_crRNA49 | GCCGCUAGAGGUGAAAUUCU |
| RN45s_crRNA50 | GGAAUAAUGGAAUAGGACCG |
| RN45s_crRNA51 | UUCGCUCGCGCUCCCUUACC |
| RN45s_crRNA52 | UUUGAGAGGCCUGGCUUUCG |
| RN45s_crRNA53 | CGUUGCAUACCCUUCCCGUC |
| RN45s_crRNA54 | UGAGCCCCUGCCGCACCCGC |
| RN45s_crRNA55 | ACCACGGGUGACGGGGAAUC |
| RN45s_crRNA56 | GCUCUUAGCUGAGUGUCCCG |
| RN45s_crRNA57 | CUAGGUGCCUGCUUCUGAGC |
| RN45s_crRNA58 | UUCGGGUUUCCCGCGCCUGC |
| RN45s_crRNA59 | UCCUCCCCGCUCGCCGCAGC |
| RN45s_crRNA60 | UGGGGAGUGAAUGGUGCUAC |
| RN45s_crRNA61 | UCUCGCGGGCGUCCGCGCGG |
| RN45s_crRNA62 | UGUGGUGGUGGCUGGGGAGA |
| RN45s_crRNA63 | CGGAAGGGCACCACCAGGAG |
| RN45s_crRNA64 | GGUAGGCAACGGUGGGCUCC |
| RN45s_crRNA65 | CCCGUUUUUGUUCCUUCCGC |
| RN45s_crRNA66 | GCCGGGGGCUGGCCGCUGUC |
| RN5.8s_crRNA67 | CGACACUUCGAACGCACUUG |
| RN5.8s_crNA68 | ACUCUUAGCGGUGGAUCACU |
| RN5.8s_crRNA69 | CGAGAAUUAAUGUGAAUUGC |

**5X Cas9 Buffer Composition**

| Reagent | Stock Concentration | Final Concentration | 50 mL Example |
|---|---|---|---|
| HEPES pH 7.4 | 1 M | 100 mM | 5 mL |
| KCl | 2 M | 500 mM | 12.5 mL |
| $MgCl_2$ | 1 M | 25 mM | 1.25 mL |
| DTT | 100 mM | 5 mM | 2.5 mL |
| Glycerol | 100 % | 25% | 12.5 mL |
| Nuclease-Free $H_20$ | | | 16.25 mL |
| **Total** | | | **50 mL** |

Supplementary table 2. Cell cycle-specific gene clusters.

| genes | cluster | biotype |
|---|---|---|
| ABCC3 | 3 | protein_coding |
| ADGRG1 | 3 | protein_coding |
| AMIGO2 | 3 | protein_coding |
| AP1M2 | 3 | protein_coding |
| ARSD | 3 | protein_coding |
| ASS1 | 3 | protein_coding |
| BCAS4 | 3 | protein_coding |
| BCL9L | 3 | protein_coding |
| C19orf33 | 3 | protein_coding |
| CA12 | 3 | protein_coding |
| CLDN4 | 3 | protein_coding |
| CLDN7 | 3 | protein_coding |
| CRABP2 | 3 | protein_coding |
| CRIP2 | 3 | protein_coding |
| CXXC5 | 3 | protein_coding |
| DKK1 | 3 | protein_coding |
| DSCAM | 3 | protein_coding |
| FAM46A | 3 | protein_coding |
| FOSL2 | 3 | protein_coding |
| GFRA1 | 3 | protein_coding |
| GPRC5A | 3 | protein_coding |
| GRHL2 | 3 | protein_coding |
| H2AFJ | 3 | protein_coding |
| HAGLROS | 3 | lincRNA |
| HSPB1 | 3 | protein_coding |
| IER3 | 3 | protein_coding |
| INPP4B | 3 | protein_coding |
| ISG15 | 3 | protein_coding |
| JUNB | 3 | protein_coding |
| KCNK6 | 3 | protein_coding |
| KRT19 | 3 | protein_coding |
| KRT80 | 3 | protein_coding |
| KYNU | 3 | protein_coding |
| LGALS3BP | 3 | protein_coding |
| LINC00052 | 3 | lincRNA |
| LINC00847 | 3 | lincRNA |
| LINC01006 | 3 | lincRNA |

| | | |
|---|---|---|
| LINC01348 | 3 | lincRNA |
| LITAF | 3 | protein_coding |
| LRP10 | 3 | protein_coding |
| NCOA3 | 3 | protein_coding |
| NPEPL1 | 3 | protein_coding |
| OCIAD2 | 3 | protein_coding |
| OLFM1 | 3 | protein_coding |
| PLK2 | 3 | protein_coding |
| PSME1 | 3 | protein_coding |
| PVRL2 | 3 | protein_coding |
| PVT1 | 3 | lincRNA |
| PYCARD | 3 | protein_coding |
| RASD1 | 3 | protein_coding |
| RBM47 | 3 | protein_coding |
| RHOBTB3 | 3 | protein_coding |
| RHOD | 3 | protein_coding |
| S100A14 | 3 | protein_coding |
| SEMA3C | 3 | protein_coding |
| SLC12A9 | 3 | protein_coding |
| SNORA28 | 3 | snoRNA |
| SNORA2B | 3 | snoRNA |
| SNORD111B | 3 | snoRNA |
| SNORD46 | 3 | snoRNA |
| SNORD59A | 3 | snoRNA |
| SNORD66 | 3 | snoRNA |
| SNORD71 | 3 | snoRNA |
| SPTSSB | 3 | protein_coding |
| ST3GAL1 | 3 | protein_coding |
| SYNPO2 | 3 | protein_coding |
| SYTL2 | 3 | protein_coding |
| TACSTD2 | 3 | protein_coding |
| TAPBP | 3 | protein_coding |
| TBC1D9 | 3 | protein_coding |
| TFF1 | 3 | protein_coding |
| TFPT | 3 | protein_coding |
| THSD4 | 3 | protein_coding |
| TRIB1 | 3 | protein_coding |
| TRPS1 | 3 | protein_coding |
| TSPYL5 | 3 | protein_coding |

| | | |
|---|---|---|
| TSTD1 | 3 | protein_coding |
| U47924.27 | 3 | NA |
| USP32 | 3 | protein_coding |
| VMP1 | 3 | protein_coding |
| WISP2 | 3 | protein_coding |
| ZNF217 | 3 | protein_coding |
| ABLIM3 | 4 | protein_coding |
| ADGRE5 | 4 | protein_coding |
| ANXA1 | 4 | protein_coding |
| ARHGAP18 | 4 | protein_coding |
| BCYRN1 | 4 | lincRNA |
| C2CD2 | 4 | protein_coding |
| CAPG | 4 | protein_coding |
| CD44 | 4 | protein_coding |
| COL6A3 | 4 | protein_coding |
| DPYSL3 | 4 | protein_coding |
| DUSP5 | 4 | protein_coding |
| EHBP1L1 | 4 | protein_coding |
| ITGA5 | 4 | protein_coding |
| KCNMA1 | 4 | protein_coding |
| KLF6 | 4 | protein_coding |
| LGALS1 | 4 | protein_coding |
| LINC00327 | 4 | lincRNA |
| LINC00702 | 4 | lincRNA |
| MIR27A | 4 | miRNA |
| MIR4435.2HG | 4 | lincRNA |
| MIRLET7D | 4 | miRNA |
| MIRLET7F1 | 4 | miRNA |
| MIRLET7F2 | 4 | miRNA |
| MME | 4 | protein_coding |
| MMP2 | 4 | protein_coding |
| MT2A | 4 | protein_coding |
| NAV2 | 4 | protein_coding |
| NCEH1 | 4 | protein_coding |
| NRP1 | 4 | protein_coding |
| PPP1R18 | 4 | protein_coding |
| PRRX2 | 4 | protein_coding |
| RIN1 | 4 | protein_coding |
| RNU1.1 | 4 | snRNA |

| | | |
|---|---|---|
| RNU1.2 | 4 | snRNA |
| RNU1.27P | 4 | snRNA |
| RNU1.28P | 4 | snRNA |
| RNVU1.18 | 4 | snRNA |
| RNVU1.7 | 4 | snRNA |
| S100A16 | 4 | protein_coding |
| S100A6 | 4 | protein_coding |
| SHCBP1 | 4 | protein_coding |
| SMIM3 | 4 | protein_coding |
| SNHG18 | 4 | lincRNA |
| SNHG23 | 4 | lincRNA |
| SQRDL | 4 | protein_coding |
| XYLT1 | 4 | protein_coding |
| ADAM12 | 5 | protein_coding |
| ADAMTS1 | 5 | protein_coding |
| ADM | 5 | protein_coding |
| ANTXR2 | 5 | protein_coding |
| ARHGAP23 | 5 | protein_coding |
| ARHGEF2 | 5 | protein_coding |
| CAV1 | 5 | protein_coding |
| CAV2 | 5 | protein_coding |
| CCBE1 | 5 | protein_coding |
| CCDC80 | 5 | protein_coding |
| CCDC88A | 5 | protein_coding |
| CDC42EP3 | 5 | protein_coding |
| CECR7 | 5 | lincRNA |
| CERKL | 5 | protein_coding |
| CLDN11 | 5 | protein_coding |
| CSPG4 | 5 | protein_coding |
| CTA.228A9.3 | 5 | NA |
| DBET | 5 | lincRNA |
| DDR2 | 5 | protein_coding |
| DKFZP434A062 | 5 | lincRNA |
| EHD2 | 5 | protein_coding |
| ENOX1.AS1 | 5 | lincRNA |
| EVA1A | 5 | protein_coding |
| FBN1 | 5 | protein_coding |
| FGF5 | 5 | protein_coding |
| FGFR1 | 5 | protein_coding |

| | | |
|---|---|---|
| FLJ26245 | 5 | lincRNA |
| FLRT2 | 5 | protein_coding |
| FP671120.6 | 5 | NA |
| FP671120.7 | 5 | NA |
| GREM1 | 5 | protein_coding |
| LINC00331 | 5 | lincRNA |
| LINC00407 | 5 | lincRNA |
| LINC00511 | 5 | lincRNA |
| LINC00836 | 5 | lincRNA |
| LINC00883 | 5 | lincRNA |
| LINC01204 | 5 | lincRNA |
| LINC01209 | 5 | lincRNA |
| LINC01297 | 5 | lincRNA |
| LINC01435 | 5 | lincRNA |
| LINC01484 | 5 | lincRNA |
| LMO7 | 5 | protein_coding |
| MEG3 | 5 | lincRNA |
| MIR181A1 | 5 | miRNA |
| MIR181A2 | 5 | miRNA |
| MIR221 | 5 | miRNA |
| MIR222 | 5 | miRNA |
| MIR222HG | 5 | lincRNA |
| MIR29A | 5 | miRNA |
| MIR31 | 5 | miRNA |
| MIR320A | 5 | miRNA |
| MIR424 | 5 | miRNA |
| MIR455 | 5 | miRNA |
| MIR615 | 5 | miRNA |
| MIR656 | 5 | miRNA |
| MIR7.2 | 5 | miRNA |
| MIR708 | 5 | miRNA |
| MSRB3 | 5 | protein_coding |
| NPTX2 | 5 | protein_coding |
| PDGFRA | 5 | protein_coding |
| PTGS1 | 5 | protein_coding |
| PTRF | 5 | protein_coding |
| RBMS2 | 5 | protein_coding |
| RCN3 | 5 | protein_coding |
| RNU1.106P | 5 | snRNA |

| | | |
|---|---|---|
| RNU1.11P | 5 | snRNA |
| RNU1.134P | 5 | snRNA |
| RNU1.42P | 5 | snRNA |
| RNU1.67P | 5 | snRNA |
| RNU12 | 5 | snRNA |
| RNU2.22P | 5 | snRNA |
| RNU2.3P | 5 | snRNA |
| RNU2.68P | 5 | snRNA |
| RNU2.69P | 5 | snRNA |
| RNU4.21P | 5 | snRNA |
| RNU4ATAC | 5 | snRNA |
| RNU5A.1 | 5 | snRNA |
| RNU5A.8P | 5 | snRNA |
| RNU5B.1 | 5 | snRNA |
| RNU5D.1 | 5 | snRNA |
| RNU5D.2P | 5 | snRNA |
| RNU5E.1 | 5 | snRNA |
| RNU5E.4P | 5 | snRNA |
| RNU6.1046P | 5 | snRNA |
| RNU6.1064P | 5 | snRNA |
| RNU6.106P | 5 | snRNA |
| RNU6.1235P | 5 | snRNA |
| RNU6.164P | 5 | snRNA |
| RNU6.30P | 5 | snRNA |
| RNU6.497P | 5 | snRNA |
| RNU6.869P | 5 | snRNA |
| RNU6ATAC | 5 | snRNA |
| RNU6ATAC25P | 5 | snRNA |
| RNU6ATAC7P | 5 | snRNA |
| RNU7.40P | 5 | snRNA |
| RNU7.93P | 5 | snRNA |
| RNVU1.15 | 5 | snRNA |
| RNVU1.17 | 5 | snRNA |
| SEC14L2 | 5 | protein_coding |
| SERPINE2 | 5 | protein_coding |
| SLC2A1.AS1 | 5 | lincRNA |
| SNORA80B | 5 | snoRNA |
| SNORD14E | 5 | snoRNA |
| SNORD93 | 5 | snoRNA |

| | | |
|---|---|---|
| SPARC | 5 | protein_coding |
| TNC | 5 | protein_coding |
| TUBA1C | 5 | protein_coding |
| TWIST2 | 5 | protein_coding |
| U8 | 5 | snoRNA |
| ZNF883 | 5 | lincRNA |
| ADAT2 | 6 | protein_coding |
| AP001172.2 | 6 | NA |
| C1orf220 | 6 | lincRNA |
| CSRP2 | 6 | protein_coding |
| CXorf57 | 6 | protein_coding |
| FAM87A | 6 | lincRNA |
| GS1.166A23.2 | 6 | NA |
| GS1.421I3.5 | 6 | NA |
| HOXB9 | 6 | protein_coding |
| HOXD13 | 6 | protein_coding |
| ID4 | 6 | protein_coding |
| LINC00467 | 6 | lincRNA |
| LINC00635 | 6 | lincRNA |
| LINC00870 | 6 | lincRNA |
| LINC00926 | 6 | lincRNA |
| LINC00971 | 6 | lincRNA |
| LINC00998 | 6 | lincRNA |
| LINC01108 | 6 | lincRNA |
| LINC01122 | 6 | lincRNA |
| LINC01605 | 6 | lincRNA |
| MARK1 | 6 | protein_coding |
| MIR320B2 | 6 | miRNA |
| MIR3687.1 | 6 | miRNA |
| MIR3928 | 6 | miRNA |
| MIR4466 | 6 | miRNA |
| MIR877 | 6 | miRNA |
| MIR941.5 | 6 | miRNA |
| NKX2.5 | 6 | protein_coding |
| PLS3 | 6 | protein_coding |
| RNF157 | 6 | protein_coding |
| RNU1.83P | 6 | snRNA |
| RNU2.17P | 6 | snRNA |
| RNU2.46P | 6 | snRNA |

| | | |
|---|---|---|
| RNU2.55P | 6 | snRNA |
| RNU6.1164P | 6 | snRNA |
| RNU6.1230P | 6 | snRNA |
| RNU6.1284P | 6 | snRNA |
| RNU6.181P | 6 | snRNA |
| RNU6.225P | 6 | snRNA |
| RNU6.27P | 6 | snRNA |
| RNU6.298P | 6 | snRNA |
| RNU6.37P | 6 | snRNA |
| RNU6.45P | 6 | snRNA |
| RNU6.489P | 6 | snRNA |
| RNU6.62P | 6 | snRNA |
| RNU6.6P | 6 | snRNA |
| RNU6.735P | 6 | snRNA |
| RNU6.738P | 6 | snRNA |
| RNU6.768P | 6 | snRNA |
| RNU6.784P | 6 | snRNA |
| RNU6.838P | 6 | snRNA |
| RNU6.883P | 6 | snRNA |
| RNU6.906P | 6 | snRNA |
| RP3.368B9.2 | 6 | NA |
| SLC6A15 | 6 | protein_coding |
| SNORA27 | 6 | snoRNA |
| TUB | 6 | protein_coding |
| ZCCHC11 | 6 | protein_coding |
| ZNF833P | 6 | lincRNA |
| ADCK3 | 7 | protein_coding |
| ANKRD18A | 7 | protein_coding |
| ARRB2 | 7 | protein_coding |
| ASRGL1 | 7 | protein_coding |
| BCOR | 7 | protein_coding |
| BEX2 | 7 | protein_coding |
| CA2 | 7 | protein_coding |
| CABLES1 | 7 | protein_coding |
| CCNE1 | 7 | protein_coding |
| CKMT1A | 7 | protein_coding |
| CKMT1B | 7 | protein_coding |
| DLX2 | 7 | protein_coding |
| DSG2 | 7 | protein_coding |

| | | |
|---|---|---|
| EPB41 | 7 | protein_coding |
| FAM157B | 7 | lincRNA |
| FGFR3 | 7 | protein_coding |
| GTPBP3 | 7 | protein_coding |
| HSPA4L | 7 | protein_coding |
| JPX | 7 | lincRNA |
| LINC00205 | 7 | lincRNA |
| MLK4 | 7 | protein_coding |
| ONECUT2 | 7 | protein_coding |
| PITX1 | 7 | protein_coding |
| RBM41 | 7 | protein_coding |
| RMND5A | 7 | protein_coding |
| RNU1.85P | 7 | snRNA |
| RRAGD | 7 | protein_coding |
| SCARNA1 | 7 | scaRNA |
| SCARNA23 | 7 | scaRNA |
| SPEN | 7 | protein_coding |
| SYNGR1 | 7 | protein_coding |
| TM2D3 | 7 | protein_coding |
| TMEM170A | 7 | protein_coding |
| ZNF704 | 7 | protein_coding |
| ZNF711 | 7 | protein_coding |
| C1orf229 | 8 | lincRNA |
| C8orf49 | 8 | lincRNA |
| COQ10A | 8 | protein_coding |
| CTC.241N9.1 | 8 | NA |
| CTC.444N24.8 | 8 | NA |
| DANT2 | 8 | lincRNA |
| FAM157C | 8 | lincRNA |
| FAM215B | 8 | lincRNA |
| FENDRR | 8 | lincRNA |
| FLVCR1.AS1 | 8 | lincRNA |
| GREB1L | 8 | protein_coding |
| HEY1 | 8 | protein_coding |
| HOXA13 | 8 | protein_coding |
| LINC00115 | 8 | lincRNA |
| LINC00630 | 8 | lincRNA |
| LINC00648 | 8 | lincRNA |
| LINC01132 | 8 | lincRNA |

| | | |
|---|---|---|
| LINC01516 | 8 | lincRNA |
| MAPKAPK5.AS1 | 8 | lincRNA |
| MIR1226 | 8 | miRNA |
| PCOLCE2 | 8 | protein_coding |
| PHF10 | 8 | protein_coding |
| POU3F2 | 8 | protein_coding |
| RAC3 | 8 | protein_coding |
| RNU1.7P | 8 | snRNA |
| RNU6.1060P | 8 | snRNA |
| RNU6.1080P | 8 | snRNA |
| RNU6.1089P | 8 | snRNA |
| RNU6.1225P | 8 | snRNA |
| RNU6.1323P | 8 | snRNA |
| RNU6.166P | 8 | snRNA |
| RNU6.16P | 8 | snRNA |
| RNU6.34P | 8 | snRNA |
| RNU6.450P | 8 | snRNA |
| RNU6.597P | 8 | snRNA |
| RNU6.923P | 8 | snRNA |
| SNORA25 | 8 | snoRNA |
| SNORA38 | 8 | snoRNA |
| SNORA59A | 8 | snoRNA |
| SNORA59B | 8 | snoRNA |
| SNORA71A | 8 | snoRNA |
| SNORA9 | 8 | snoRNA |
| SNORD101 | 8 | snoRNA |
| SNORD117 | 8 | snoRNA |
| SNORD127 | 8 | snoRNA |
| SNORD17 | 8 | snoRNA |
| SNORD1B | 8 | snoRNA |
| SNORD62A | 8 | snoRNA |
| SNORD92 | 8 | snoRNA |
| SRP14.AS1 | 8 | lincRNA |
| URB2 | 8 | protein_coding |
| ADGRG6 | 9 | protein_coding |
| AGR2 | 9 | protein_coding |
| AGRN | 9 | protein_coding |
| ATP2A3 | 9 | protein_coding |
| C1QTNF6 | 9 | protein_coding |

| | | |
|---|---|---|
| CACNG4 | 9 | protein_coding |
| CD24 | 9 | protein_coding |
| CLDN3 | 9 | protein_coding |
| CLU | 9 | protein_coding |
| CTC.236F12.4 | 9 | NA |
| CYBA | 9 | protein_coding |
| EPCAM | 9 | protein_coding |
| ESRP1 | 9 | protein_coding |
| FAM174B | 9 | protein_coding |
| G6PD | 9 | protein_coding |
| GATA3.AS1 | 9 | lincRNA |
| KCNK15 | 9 | protein_coding |
| LINC00623 | 9 | lincRNA |
| LINC01186 | 9 | lincRNA |
| LXN | 9 | protein_coding |
| LYPD3 | 9 | protein_coding |
| MAL2 | 9 | protein_coding |
| MIR141 | 9 | miRNA |
| MIR200C | 9 | miRNA |
| MIR3929 | 9 | miRNA |
| MIR4737 | 9 | miRNA |
| N4BP3 | 9 | protein_coding |
| NRCAM | 9 | protein_coding |
| PPAP2C | 9 | protein_coding |
| PPP1R16A | 9 | protein_coding |
| SEMA3B | 9 | protein_coding |
| SEMA4B | 9 | protein_coding |
| SIK1 | 9 | protein_coding |
| SLC7A2 | 9 | protein_coding |
| SLC9A3R1 | 9 | protein_coding |
| SNORD94 | 9 | snoRNA |
| SPINT1 | 9 | protein_coding |
| TMEM64 | 9 | protein_coding |
| TSPAN15 | 9 | protein_coding |
| U2 | 9 | snRNA |
| AEBP1 | 10 | protein_coding |
| AF131217.1 | 10 | NA |
| AP001476.3 | 10 | NA |
| AP001607.1 | 10 | NA |

| | | |
|---|---|---|
| AXL | 10 | protein_coding |
| CASP4 | 10 | protein_coding |
| CLEC2B | 10 | protein_coding |
| CPED1 | 10 | protein_coding |
| CTC.384G19.1 | 10 | NA |
| CYB5R2 | 10 | protein_coding |
| EMP1 | 10 | protein_coding |
| ENPP2 | 10 | protein_coding |
| FAM138D | 10 | lincRNA |
| FOSL1 | 10 | protein_coding |
| FST | 10 | protein_coding |
| GAPLINC | 10 | lincRNA |
| GLIPR1 | 10 | protein_coding |
| HAS2 | 10 | protein_coding |
| IFI16 | 10 | protein_coding |
| LINC00346 | 10 | lincRNA |
| LINC00565 | 10 | lincRNA |
| LINC00667 | 10 | lincRNA |
| LINC00704 | 10 | lincRNA |
| LINC00707 | 10 | lincRNA |
| LINC00941 | 10 | lincRNA |
| LINC01095 | 10 | lincRNA |
| LINC01133 | 10 | lincRNA |
| LINC01230 | 10 | lincRNA |
| LTBP1 | 10 | protein_coding |
| LY6K | 10 | protein_coding |
| MEG8 | 10 | lincRNA |
| MIR100 | 10 | miRNA |
| MIR100HG | 10 | lincRNA |
| MIR125B1 | 10 | miRNA |
| MIR130A | 10 | miRNA |
| MIR137HG | 10 | lincRNA |
| MIR138.1 | 10 | miRNA |
| MIR138.2 | 10 | miRNA |
| MIR155HG | 10 | lincRNA |
| MIR193A | 10 | miRNA |
| MIR199A1 | 10 | miRNA |
| MIR199A2 | 10 | miRNA |
| MIR199B | 10 | miRNA |

| | | |
|---|---|---|
| MIR299 | 10 | miRNA |
| MIR323A | 10 | miRNA |
| MIR329.2 | 10 | miRNA |
| MIR337 | 10 | miRNA |
| MIR365B | 10 | miRNA |
| MIR369 | 10 | miRNA |
| MIR370 | 10 | miRNA |
| MIR376A1 | 10 | miRNA |
| MIR376B | 10 | miRNA |
| MIR376C | 10 | miRNA |
| MIR377 | 10 | miRNA |
| MIR379 | 10 | miRNA |
| MIR382 | 10 | miRNA |
| MIR409 | 10 | miRNA |
| MIR411 | 10 | miRNA |
| MIR412 | 10 | miRNA |
| MIR485 | 10 | miRNA |
| MIR487A | 10 | miRNA |
| MIR495 | 10 | miRNA |
| MIRLET7A2 | 10 | miRNA |
| MT1A | 10 | protein_coding |
| MT1E | 10 | protein_coding |
| NNMT | 10 | protein_coding |
| PLAU | 10 | protein_coding |
| PLAUR | 10 | protein_coding |
| PRKCDBP | 10 | protein_coding |
| PWAR6 | 10 | lincRNA |
| RFTN1 | 10 | protein_coding |
| RNU1.59P | 10 | snRNA |
| RNU1.84P | 10 | snRNA |
| RP6.109B7.2 | 10 | NA |
| RP6.109B7.4 | 10 | NA |
| RPS4Y1 | 10 | protein_coding |
| SERPINE1 | 10 | protein_coding |
| SFRP1 | 10 | protein_coding |
| SH2B3 | 10 | protein_coding |
| SLC17A9 | 10 | protein_coding |
| SLC43A3 | 10 | protein_coding |
| SNHG24 | 10 | lincRNA |

| | | |
|---|---|---|
| SNORA48 | 10 | snoRNA |
| SNORD113 | 10 | snoRNA |
| SNORD113.3 | 10 | snoRNA |
| SNORD113.4 | 10 | snoRNA |
| SNORD113.5 | 10 | snoRNA |
| SNORD113.6 | 10 | snoRNA |
| SNORD113.7 | 10 | snoRNA |
| SNORD113.8 | 10 | snoRNA |
| SNORD113.9 | 10 | snoRNA |
| SNORD114.1 | 10 | snoRNA |
| SNORD114.10 | 10 | snoRNA |
| SNORD114.11 | 10 | snoRNA |
| SNORD114.12 | 10 | snoRNA |
| SNORD114.13 | 10 | snoRNA |
| SNORD114.14 | 10 | snoRNA |
| SNORD114.15 | 10 | snoRNA |
| SNORD114.17 | 10 | snoRNA |
| SNORD114.20 | 10 | snoRNA |
| SNORD114.21 | 10 | snoRNA |
| SNORD114.22 | 10 | snoRNA |
| SNORD114.23 | 10 | snoRNA |
| SNORD114.25 | 10 | snoRNA |
| SNORD114.28 | 10 | snoRNA |
| SNORD114.3 | 10 | snoRNA |
| SNORD114.6 | 10 | snoRNA |
| SNORD114.9 | 10 | snoRNA |
| SNORD116.1 | 10 | snoRNA |
| SNORD116.13 | 10 | snoRNA |
| SNORD116.14 | 10 | snoRNA |
| SNORD116.15 | 10 | snoRNA |
| SNORD116.16 | 10 | snoRNA |
| SNORD116.17 | 10 | snoRNA |
| SNORD116.19 | 10 | snoRNA |
| SNORD116.23 | 10 | snoRNA |
| SNORD116.24 | 10 | snoRNA |
| SNORD116.25 | 10 | snoRNA |
| SNORD116.3 | 10 | snoRNA |
| SNORD116.5 | 10 | snoRNA |
| SNORD116.6 | 10 | snoRNA |

| | | |
|---|---|---|
| SNORD116.7 | 10 | snoRNA |
| SNORD116.8 | 10 | snoRNA |
| SNORD116.9 | 10 | snoRNA |
| SNORD123 | 10 | snoRNA |
| SNORD48 | 10 | snoRNA |
| TMBIM1 | 10 | protein_coding |
| TTTY14 | 10 | lincRNA |
| TTTY15 | 10 | lincRNA |
| WI2.87327B8.1 | 10 | NA |
| AF003625.3 | 11 | NA |
| AP000472.2 | 11 | NA |
| AP000704.5 | 11 | NA |
| BARX1.AS1 | 11 | lincRNA |
| BSN.AS2 | 11 | lincRNA |
| BX255923.3 | 11 | NA |
| CEP83.AS1 | 11 | lincRNA |
| CTC.459F4.3 | 11 | NA |
| FAM66B | 11 | lincRNA |
| FAM87B | 11 | lincRNA |
| GHET1 | 11 | lincRNA |
| HOXA9 | 11 | protein_coding |
| KB.1471A8.1 | 11 | NA |
| KIAA0087 | 11 | lincRNA |
| LINC00106 | 11 | lincRNA |
| LINC00282 | 11 | lincRNA |
| LINC00294 | 11 | lincRNA |
| LINC00324 | 11 | lincRNA |
| LINC00441 | 11 | lincRNA |
| LINC00574 | 11 | lincRNA |
| LINC00939 | 11 | lincRNA |
| LINC01010 | 11 | lincRNA |
| LINC01018 | 11 | lincRNA |
| LINC01144 | 11 | lincRNA |
| LINC01351 | 11 | lincRNA |
| LINC01460 | 11 | lincRNA |
| LINC01560 | 11 | lincRNA |
| MID1 | 11 | protein_coding |
| MIR1224 | 11 | miRNA |
| MIR30B | 11 | miRNA |

| | | |
|---|---|---|
| MIR374A | 11 | miRNA |
| MIR3945 | 11 | miRNA |
| MIR503 | 11 | miRNA |
| MIR542 | 11 | miRNA |
| NAMA | 11 | lincRNA |
| OGFRP1 | 11 | lincRNA |
| RAMP2.AS1 | 11 | lincRNA |
| RNU1.103P | 11 | snRNA |
| RNU1.18P | 11 | snRNA |
| RNU1.72P | 11 | snRNA |
| RNU2.56P | 11 | snRNA |
| RNU4.40P | 11 | snRNA |
| RNU6.1099P | 11 | snRNA |
| RNU6.1103P | 11 | snRNA |
| RNU6.1104P | 11 | snRNA |
| RNU6.1122P | 11 | snRNA |
| RNU6.114P | 11 | snRNA |
| RNU6.1195P | 11 | snRNA |
| RNU6.119P | 11 | snRNA |
| RNU6.1264P | 11 | snRNA |
| RNU6.1301P | 11 | snRNA |
| RNU6.140P | 11 | snRNA |
| RNU6.14P | 11 | snRNA |
| RNU6.18P | 11 | snRNA |
| RNU6.196P | 11 | snRNA |
| RNU6.204P | 11 | snRNA |
| RNU6.28P | 11 | snRNA |
| RNU6.31P | 11 | snRNA |
| RNU6.322P | 11 | snRNA |
| RNU6.33P | 11 | snRNA |
| RNU6.393P | 11 | snRNA |
| RNU6.3P | 11 | snRNA |
| RNU6.41P | 11 | snRNA |
| RNU6.429P | 11 | snRNA |
| RNU6.43P | 11 | snRNA |
| RNU6.46P | 11 | snRNA |
| RNU6.481P | 11 | snRNA |
| RNU6.487P | 11 | snRNA |
| RNU6.490P | 11 | snRNA |

| | | |
|---|---|---|
| RNU6.498P | 11 | snRNA |
| RNU6.4P | 11 | snRNA |
| RNU6.500P | 11 | snRNA |
| RNU6.5P | 11 | snRNA |
| RNU6.637P | 11 | snRNA |
| RNU6.702P | 11 | snRNA |
| RNU6.756P | 11 | snRNA |
| RNU6.761P | 11 | snRNA |
| RNU6.79P | 11 | snRNA |
| RNU6.842P | 11 | snRNA |
| RNU6.854P | 11 | snRNA |
| RNU6.9 | 11 | snRNA |
| RNU6.950P | 11 | snRNA |
| RNU6.97P | 11 | snRNA |
| RNU7.41P | 11 | snRNA |
| RP3.466P17.1 | 11 | NA |
| SH3RF3.AS1 | 11 | lincRNA |
| SLC25A30.AS1 | 11 | lincRNA |
| SNORD124 | 11 | snoRNA |
| SNORD7 | 11 | snoRNA |
| Z83851.4 | 11 | NA |
| AF038458.5 | 12 | NA |
| AP003900.6 | 12 | NA |
| B4GAT1 | 12 | protein_coding |
| BEX4 | 12 | protein_coding |
| C21orf62.AS1 | 12 | lincRNA |
| CDKN2A | 12 | protein_coding |
| DPYSL5 | 12 | protein_coding |
| FAM184A | 12 | protein_coding |
| FAM27C | 12 | lincRNA |
| FBN2 | 12 | protein_coding |
| HOXA10 | 12 | protein_coding |
| INA | 12 | protein_coding |
| IRS4 | 12 | protein_coding |
| KC6 | 12 | lincRNA |
| LIN28B | 12 | protein_coding |
| LINC00240 | 12 | lincRNA |
| LINC00472 | 12 | lincRNA |
| LINC00491 | 12 | lincRNA |

| | | |
|---|---|---|
| LINC00543 | 12 | lincRNA |
| LINC00703 | 12 | lincRNA |
| LINC01422 | 12 | lincRNA |
| LOH12CR2 | 12 | lincRNA |
| MIR101.1 | 12 | miRNA |
| MIR218.1 | 12 | miRNA |
| PAUPAR | 12 | lincRNA |
| PYGL | 12 | protein_coding |
| RNU1.115P | 12 | snRNA |
| RNU1.65P | 12 | snRNA |
| RNU5F.1 | 12 | snRNA |
| RNU6.1005P | 12 | snRNA |
| RNU6.1038P | 12 | snRNA |
| RNU6.1124P | 12 | snRNA |
| RNU6.116P | 12 | snRNA |
| RNU6.1201P | 12 | snRNA |
| RNU6.1255P | 12 | snRNA |
| RNU6.128P | 12 | snRNA |
| RNU6.12P | 12 | snRNA |
| RNU6.1329P | 12 | snRNA |
| RNU6.13P | 12 | snRNA |
| RNU6.154P | 12 | snRNA |
| RNU6.15P | 12 | snRNA |
| RNU6.216P | 12 | snRNA |
| RNU6.21P | 12 | snRNA |
| RNU6.222P | 12 | snRNA |
| RNU6.227P | 12 | snRNA |
| RNU6.228P | 12 | snRNA |
| RNU6.22P | 12 | snRNA |
| RNU6.238P | 12 | snRNA |
| RNU6.32P | 12 | snRNA |
| RNU6.359P | 12 | snRNA |
| RNU6.378P | 12 | snRNA |
| RNU6.40P | 12 | snRNA |
| RNU6.453P | 12 | snRNA |
| RNU6.460P | 12 | snRNA |
| RNU6.463P | 12 | snRNA |
| RNU6.496P | 12 | snRNA |
| RNU6.510P | 12 | snRNA |

| | | |
|---|---|---|
| RNU6.533P | 12 | snRNA |
| RNU6.574P | 12 | snRNA |
| RNU6.578P | 12 | snRNA |
| RNU6.7 | 12 | snRNA |
| RNU6.729P | 12 | snRNA |
| RNU6.737P | 12 | snRNA |
| RNU6.73P | 12 | snRNA |
| RNU6.759P | 12 | snRNA |
| RNU6.905P | 12 | snRNA |
| RNU6.937P | 12 | snRNA |
| RNU6.996P | 12 | snRNA |
| RNU6ATAC28P | 12 | snRNA |
| RNU6ATAC37P | 12 | snRNA |
| RP3.388N13.5 | 12 | NA |
| RP3.460G2.2 | 12 | NA |
| SCARNA12 | 12 | snoRNA |
| SLITRK5 | 12 | protein_coding |
| SNORA11 | 12 | snoRNA |
| SNORA11C | 12 | snoRNA |
| SNORA72 | 12 | snoRNA |
| SNORD3A | 12 | snoRNA |
| U6 | 12 | snRNA |
| ZNF280B | 12 | protein_coding |
| AF064858.8 | 13 | NA |
| ARHGAP19 | 13 | protein_coding |
| BUB1 | 13 | protein_coding |
| BUB1B | 13 | protein_coding |
| CCNA2 | 13 | protein_coding |
| CDC25C | 13 | protein_coding |
| CENPE | 13 | protein_coding |
| CENPF | 13 | protein_coding |
| CKAP2 | 13 | protein_coding |
| CKAP5 | 13 | protein_coding |
| CNTRL | 13 | protein_coding |
| CTC.756D1.3 | 13 | NA |
| CYP4F35P | 13 | lincRNA |
| DEPDC1 | 13 | protein_coding |
| ERCC6L | 13 | protein_coding |
| FOCAD.AS1 | 13 | lincRNA |

| | | |
|---|---|---|
| GAS2L3 | 13 | protein_coding |
| GTSE1 | 13 | protein_coding |
| HMGB2 | 13 | protein_coding |
| KIF11 | 13 | protein_coding |
| KIF14 | 13 | protein_coding |
| KIF15 | 13 | protein_coding |
| KIF18A | 13 | protein_coding |
| KIF18B | 13 | protein_coding |
| KIF20B | 13 | protein_coding |
| KIF4A | 13 | protein_coding |
| LINC00299 | 13 | lincRNA |
| LINC01579 | 13 | lincRNA |
| MIR15A | 13 | miRNA |
| MIR26B | 13 | miRNA |
| MIR550A1 | 13 | miRNA |
| MIR550A2 | 13 | miRNA |
| MIR941.4 | 13 | miRNA |
| NCAPG | 13 | protein_coding |
| NEK2 | 13 | protein_coding |
| NUF2 | 13 | protein_coding |
| PLK1 | 13 | protein_coding |
| RNU6.1077P | 13 | snRNA |
| RNU6.1188P | 13 | snRNA |
| RNU6.484P | 13 | snRNA |
| RNU6.70P | 13 | snRNA |
| RNU6.748P | 13 | snRNA |
| RNU6ATAC3P | 13 | snRNA |
| RNU7.79P | 13 | snRNA |
| RP3.510D11.1 | 13 | NA |
| SCGB1B2P | 13 | lincRNA |
| SPAG5 | 13 | protein_coding |
| SPC25 | 13 | protein_coding |
| TOP2A | 13 | protein_coding |
| TROAP | 13 | protein_coding |
| TTK | 13 | protein_coding |
| UBE2S | 13 | protein_coding |
| USP27X.AS1 | 13 | lincRNA |
| AF127936.7 | 14 | NA |
| AP000253.1 | 14 | NA |

| | | |
|---|---|---|
| AP006222.2 | 14 | NA |
| CLIP1.AS1 | 14 | lincRNA |
| HES4 | 14 | protein_coding |
| HRAT92 | 14 | lincRNA |
| LINC00618 | 14 | lincRNA |
| LINC00680 | 14 | lincRNA |
| LINC00997 | 14 | lincRNA |
| LINC01123 | 14 | lincRNA |
| LINC01184 | 14 | lincRNA |
| LINC01355 | 14 | lincRNA |
| LINC01521 | 14 | lincRNA |
| LINC01547 | 14 | lincRNA |
| MIR3180.4 | 14 | miRNA |
| MIR6765 | 14 | miRNA |
| MNX1.AS1 | 14 | lincRNA |
| RNASET2 | 14 | protein_coding |
| RNU2.61P | 14 | snRNA |
| RNU6.209P | 14 | snRNA |
| SCARNA14 | 14 | scaRNA |
| SNORD2 | 14 | snoRNA |
| SNORD91A | 14 | snoRNA |
| AP006621.5 | 16 | NA |
| ARL17A | 16 | protein_coding |
| ARL17B | 16 | protein_coding |
| CACHD1 | 16 | protein_coding |
| CENPV | 16 | protein_coding |
| CKB | 16 | protein_coding |
| CTC.492K19.7 | 16 | NA |
| DGCR8 | 16 | protein_coding |
| DUSP22 | 16 | protein_coding |
| FLVCR1 | 16 | protein_coding |
| FOXO1 | 16 | protein_coding |
| MAF | 16 | protein_coding |
| RNU6.1108P | 16 | snRNA |
| RNU6.1321P | 16 | snRNA |
| RNU6.35P | 16 | snRNA |
| RNU6.576P | 16 | snRNA |
| RNU6.660P | 16 | snRNA |
| RNU6.722P | 16 | snRNA |

| | | |
|---|---|---|
| RNU6.806P | 16 | snRNA |
| RNU6.890P | 16 | snRNA |
| RNU6.891P | 16 | snRNA |
| RNU6.959P | 16 | snRNA |
| RNU6V | 16 | snRNA |
| AIFM1 | 17 | protein_coding |
| CASC9 | 17 | lincRNA |
| CDC37L1.AS1 | 17 | lincRNA |
| CTC.444N24.11 | 17 | NA |
| DACH1 | 17 | protein_coding |
| DKFZP434L187 | 17 | lincRNA |
| EN2 | 17 | protein_coding |
| GAL | 17 | protein_coding |
| GLIDR | 17 | lincRNA |
| HS6ST2 | 17 | protein_coding |
| LINC01003 | 17 | lincRNA |
| LL0XNC01.237H1.2 | 17 | NA |
| LL22NC03.2H8.5 | 17 | NA |
| RN7SL832P | 17 | lincRNA |
| RNU6.8 | 17 | snRNA |
| SCARNA10 | 17 | snoRNA |
| SLC25A23 | 17 | protein_coding |
| SNORA47 | 17 | snoRNA |
| SNORA79 | 17 | snoRNA |
| SNORD11B | 17 | snoRNA |
| TXNDC16 | 17 | protein_coding |
| AJ011932.1 | 19 | NA |
| C20orf197 | 19 | lincRNA |
| CTC.529P8.1 | 19 | NA |
| KB.1440D3.14 | 19 | NA |
| KB.1980E6.3 | 19 | NA |
| LINC01060 | 19 | lincRNA |
| LINC01114 | 19 | lincRNA |
| LINC01444 | 19 | lincRNA |
| MIR1185.1 | 19 | miRNA |
| MIR1185.2 | 19 | miRNA |
| MIR143 | 19 | miRNA |
| MIR154 | 19 | miRNA |
| MIR30E | 19 | miRNA |

| | | |
|---|---|---|
| MIR323B | 19 | miRNA |
| MIR381 | 19 | miRNA |
| MIR487B | 19 | miRNA |
| MIR496 | 19 | miRNA |
| MIR758 | 19 | miRNA |
| RNU1.78P | 19 | snRNA |
| RP3.471M13.2 | 19 | NA |
| SNORD116.2 | 19 | snoRNA |
| SNORD116.27 | 19 | snoRNA |
| SNORD116.29 | 19 | snoRNA |
| SPRY4.IT1 | 19 | lincRNA |
| AP000230.1 | 22 | NA |
| FBXL20 | 22 | protein_coding |
| KB.1836B5.4 | 22 | NA |
| LINC00351 | 22 | lincRNA |
| LINC00379 | 22 | lincRNA |
| LINC00426 | 22 | lincRNA |
| MIR25 | 22 | miRNA |
| MIR3143 | 22 | miRNA |
| MIR365A | 22 | miRNA |
| MIR744 | 22 | miRNA |
| MIR7846 | 22 | miRNA |
| MIR99AHG | 22 | lincRNA |
| NBAT1 | 22 | lincRNA |
| RNU4ATAC18P | 22 | snRNA |
| RNU6.234P | 22 | snRNA |
| RNU6.998P | 22 | snRNA |
| AP000255.6 | 23 | NA |
| AP005530.1 | 23 | NA |
| ARSK | 23 | protein_coding |
| BCL11A | 23 | protein_coding |
| BEX1 | 23 | protein_coding |
| CCDC181 | 23 | protein_coding |
| CCDC8 | 23 | protein_coding |
| CCND2 | 23 | protein_coding |
| CDH2 | 23 | protein_coding |
| CTA.246H3.12 | 23 | NA |
| CTC.350I8.1 | 23 | NA |
| CTC.360G5.9 | 23 | NA |

| | | |
|---|---|---|
| CTC.461F20.1 | 23 | NA |
| CTC.480C2.1 | 23 | NA |
| CXADR | 23 | protein_coding |
| CYP4A22.AS1 | 23 | lincRNA |
| D21S2088E | 23 | lincRNA |
| DNAJC3.AS1 | 23 | lincRNA |
| EPHA5.AS1 | 23 | lincRNA |
| FALEC | 23 | lincRNA |
| FHL1 | 23 | protein_coding |
| FLJ12825 | 23 | lincRNA |
| FOXCUT | 23 | lincRNA |
| FOXG1 | 23 | protein_coding |
| FP671120.1 | 23 | NA |
| GATA6.AS1 | 23 | lincRNA |
| GS1.259H13.2 | 23 | NA |
| HCG11 | 23 | lincRNA |
| HHLA3 | 23 | protein_coding |
| IGF2BP1 | 23 | protein_coding |
| KBTBD11.OT1 | 23 | lincRNA |
| LINC00208 | 23 | lincRNA |
| LINC00242 | 23 | lincRNA |
| LINC00404 | 23 | lincRNA |
| LINC00937 | 23 | lincRNA |
| LINC01033 | 23 | lincRNA |
| LINC01166 | 23 | lincRNA |
| LINC01176 | 23 | lincRNA |
| LINC01234 | 23 | lincRNA |
| LINC01273 | 23 | lincRNA |
| LINC01354 | 23 | lincRNA |
| LINC01456 | 23 | lincRNA |
| LINC01535 | 23 | lincRNA |
| LINC01551 | 23 | lincRNA |
| MAP7D2 | 23 | protein_coding |
| MIR1180 | 23 | miRNA |
| MIR1277 | 23 | miRNA |
| MIR1301 | 23 | miRNA |
| MIR15B | 23 | miRNA |
| MIR196A2 | 23 | miRNA |
| MIR19B2 | 23 | miRNA |

| | | |
|---|---|---|
| MIR301B | 23 | miRNA |
| MIR3178 | 23 | miRNA |
| MIR328 | 23 | miRNA |
| MIR3685 | 23 | miRNA |
| MIR3939 | 23 | miRNA |
| MIR4470 | 23 | miRNA |
| MIR4500HG | 23 | lincRNA |
| MIR503HG | 23 | lincRNA |
| MIR551B | 23 | miRNA |
| MIR6068 | 23 | miRNA |
| MIR766 | 23 | miRNA |
| MIR92A2 | 23 | miRNA |
| NEFL | 23 | protein_coding |
| NPRL2 | 23 | protein_coding |
| PCDH9 | 23 | protein_coding |
| POU4F1 | 23 | protein_coding |
| RNF138 | 23 | protein_coding |
| RNU1.13P | 23 | snRNA |
| RNU1.88P | 23 | snRNA |
| RNU2.13P | 23 | snRNA |
| RNU2.32P | 23 | snRNA |
| RNU6.1 | 23 | snRNA |
| RNU6.1020P | 23 | snRNA |
| RNU6.1067P | 23 | snRNA |
| RNU6.1079P | 23 | snRNA |
| RNU6.1084P | 23 | snRNA |
| RNU6.10P | 23 | snRNA |
| RNU6.1140P | 23 | snRNA |
| RNU6.1141P | 23 | snRNA |
| RNU6.1145P | 23 | snRNA |
| RNU6.1159P | 23 | snRNA |
| RNU6.115P | 23 | snRNA |
| RNU6.1183P | 23 | snRNA |
| RNU6.1227P | 23 | snRNA |
| RNU6.1238P | 23 | snRNA |
| RNU6.1248P | 23 | snRNA |
| RNU6.1316P | 23 | snRNA |
| RNU6.1336P | 23 | snRNA |
| RNU6.152P | 23 | snRNA |

| | | |
|---|---|---|
| RNU6.17P | 23 | snRNA |
| RNU6.188P | 23 | snRNA |
| RNU6.20P | 23 | snRNA |
| RNU6.235P | 23 | snRNA |
| RNU6.23P | 23 | snRNA |
| RNU6.25P | 23 | snRNA |
| RNU6.263P | 23 | snRNA |
| RNU6.312P | 23 | snRNA |
| RNU6.326P | 23 | snRNA |
| RNU6.36P | 23 | snRNA |
| RNU6.380P | 23 | snRNA |
| RNU6.38P | 23 | snRNA |
| RNU6.391P | 23 | snRNA |
| RNU6.39P | 23 | snRNA |
| RNU6.428P | 23 | snRNA |
| RNU6.42P | 23 | snRNA |
| RNU6.434P | 23 | snRNA |
| RNU6.439P | 23 | snRNA |
| RNU6.44P | 23 | snRNA |
| RNU6.476P | 23 | snRNA |
| RNU6.47P | 23 | snRNA |
| RNU6.480P | 23 | snRNA |
| RNU6.49P | 23 | snRNA |
| RNU6.505P | 23 | snRNA |
| RNU6.517P | 23 | snRNA |
| RNU6.537P | 23 | snRNA |
| RNU6.553P | 23 | snRNA |
| RNU6.582P | 23 | snRNA |
| RNU6.609P | 23 | snRNA |
| RNU6.60P | 23 | snRNA |
| RNU6.629P | 23 | snRNA |
| RNU6.647P | 23 | snRNA |
| RNU6.669P | 23 | snRNA |
| RNU6.707P | 23 | snRNA |
| RNU6.723P | 23 | snRNA |
| RNU6.743P | 23 | snRNA |
| RNU6.820P | 23 | snRNA |
| RNU6.826P | 23 | snRNA |
| RNU6.857P | 23 | snRNA |

| | | |
|---|---|---|
| RNU6.862P | 23 | snRNA |
| RNU6.86P | 23 | snRNA |
| RNU6.876P | 23 | snRNA |
| RNU6.880P | 23 | snRNA |
| RNU6.91P | 23 | snRNA |
| RNU6.95P | 23 | snRNA |
| RNU6.976P | 23 | snRNA |
| RNU6.984P | 23 | snRNA |
| RNU6.994P | 23 | snRNA |
| RNU6ATAC16P | 23 | snRNA |
| RNU6ATAC26P | 23 | snRNA |
| RNU6ATAC2P | 23 | snRNA |
| RNVU1.2 | 23 | snRNA |
| RNVU1.4 | 23 | snRNA |
| SALRNA1 | 23 | lincRNA |
| SLC1A3 | 23 | protein_coding |
| SMCO4 | 23 | protein_coding |
| SNORA35 | 23 | snoRNA |
| SNORD11 | 23 | snoRNA |
| SNORD14A | 23 | snoRNA |
| SNORD69 | 23 | snoRNA |
| SNORD83A | 23 | snoRNA |
| snoU109 | 23 | snoRNA |
| TMSB15A | 23 | protein_coding |
| TRI.TAT2.3 | 23 | miRNA |
| TRO | 23 | protein_coding |
| TSIX | 23 | lincRNA |
| XYLB | 23 | protein_coding |
| ZNF582.AS1 | 23 | lincRNA |
| ZNF589 | 23 | protein_coding |
| ZNHIT6 | 23 | protein_coding |
| AP000569.9 | 26 | NA |
| C22orf34 | 26 | lincRNA |
| CTC.786C10.1 | 26 | NA |
| FAM138A | 26 | lincRNA |
| FAM138E | 26 | lincRNA |
| GS1.279B7.1 | 26 | NA |
| LINC00355 | 26 | lincRNA |
| LINC00486 | 26 | lincRNA |

| | | |
|---|---|---|
| LINC00595 | 26 | lincRNA |
| LINC01149 | 26 | lincRNA |
| LINC01405 | 26 | lincRNA |
| LINC01440 | 26 | lincRNA |
| LINC01563 | 26 | lincRNA |
| MIR329.1 | 26 | miRNA |
| MIR3943 | 26 | miRNA |
| MIR4792 | 26 | miRNA |
| MIR501 | 26 | miRNA |
| OTX2.AS1 | 26 | lincRNA |
| PGM5.AS1 | 26 | lincRNA |
| RNU6.897P | 26 | snRNA |
| TPRG1.AS1 | 26 | lincRNA |
| U6atac | 26 | snRNA |
| AP000688.29 | 27 | NA |
| CTC.537E7.3 | 27 | NA |
| LINC00484 | 27 | lincRNA |
| LINC01241 | 27 | lincRNA |
| LINC01336 | 27 | lincRNA |
| LINC01358 | 27 | lincRNA |
| MIR6871 | 27 | miRNA |
| MIR7515HG | 27 | lincRNA |
| PSMB8.AS1 | 27 | lincRNA |
| PTGES2.AS1 | 27 | lincRNA |
| RNU4.52P | 27 | snRNA |
| RNU6.571P | 27 | snRNA |
| SMC2.AS1 | 27 | lincRNA |
| SNORD109B | 27 | snoRNA |
| SNORD116.11 | 27 | snoRNA |
| SNORD116.12 | 27 | snoRNA |
| AP001628.6 | 29 | NA |
| BAIAP2.AS1 | 29 | lincRNA |
| CTA.276F8.1 | 29 | NA |
| CYP4F26P | 29 | lincRNA |
| DKFZP434I0714 | 29 | lincRNA |
| DLX2.AS1 | 29 | lincRNA |
| ERVH48.1 | 29 | lincRNA |
| KDM6B | 29 | protein_coding |
| LINC00176 | 29 | lincRNA |

| | | |
|---|---|---|
| LINC00243 | 29 | lincRNA |
| LINC00461 | 29 | lincRNA |
| LINC00562 | 29 | lincRNA |
| LINC01012 | 29 | lincRNA |
| LINC01569 | 29 | lincRNA |
| MIATNB | 29 | lincRNA |
| MIR210HG | 29 | lincRNA |
| MIR421 | 29 | miRNA |
| RP3.510D11.2 | 29 | NA |
| RP6.65G23.3 | 29 | NA |
| SCARNA5 | 29 | scaRNA |
| SNORD63 | 29 | snoRNA |
| APPBP2 | 30 | protein_coding |
| BAMBI | 30 | protein_coding |
| BCAS3 | 30 | protein_coding |
| COX6C | 30 | protein_coding |
| CTSD | 30 | protein_coding |
| DUSP16 | 30 | protein_coding |
| EPHB4 | 30 | protein_coding |
| EPPK1 | 30 | protein_coding |
| ESR1 | 30 | protein_coding |
| FAM213A | 30 | protein_coding |
| FAM83H | 30 | protein_coding |
| FAM83H.AS1 | 30 | lincRNA |
| FOXA1 | 30 | protein_coding |
| GATA3 | 30 | protein_coding |
| IDH2 | 30 | protein_coding |
| ITGB4 | 30 | protein_coding |
| JARID2 | 30 | protein_coding |
| JUP | 30 | protein_coding |
| KRT18 | 30 | protein_coding |
| KRT8 | 30 | protein_coding |
| LLGL2 | 30 | protein_coding |
| LSR | 30 | protein_coding |
| MARVELD2 | 30 | protein_coding |
| MREG | 30 | protein_coding |
| MTL5 | 30 | protein_coding |
| MYLIP | 30 | protein_coding |
| MYO5C | 30 | protein_coding |

| | | |
|---|---|---|
| NOTCH3 | 30 | protein_coding |
| PARD6B | 30 | protein_coding |
| PFDN4 | 30 | protein_coding |
| PPP1R13B | 30 | protein_coding |
| PSMD6 | 30 | protein_coding |
| SLC39A6 | 30 | protein_coding |
| SULF2 | 30 | protein_coding |
| TOB1 | 30 | protein_coding |
| TUBD1 | 30 | protein_coding |
| ARFGEF3 | 31 | protein_coding |
| ATP5EP2 | 31 | protein_coding |
| BMP7 | 31 | protein_coding |
| C17orf82 | 31 | lincRNA |
| CYP1B1 | 31 | protein_coding |
| EEF1A2 | 31 | protein_coding |
| ERBB3 | 31 | protein_coding |
| FAM84B | 31 | protein_coding |
| GREB1 | 31 | protein_coding |
| HID1 | 31 | protein_coding |
| LAMA5 | 31 | protein_coding |
| LINC00992 | 31 | lincRNA |
| RPS6KB1 | 31 | protein_coding |
| SPINT2 | 31 | protein_coding |
| STARD10 | 31 | protein_coding |
| THEM6 | 31 | protein_coding |
| TMEM189 | 31 | protein_coding |
| TPD52L1 | 31 | protein_coding |
| TRIM37 | 31 | protein_coding |
| TTC39A | 31 | protein_coding |
| VAMP8 | 31 | protein_coding |
| ARHGAP11A | 32 | protein_coding |
| ARHGAP11B | 32 | protein_coding |
| ASPM | 32 | protein_coding |
| CASC5 | 32 | protein_coding |
| CBR3 | 32 | protein_coding |
| CDCA2 | 32 | protein_coding |
| CDCA3 | 32 | protein_coding |
| CDCA8 | 32 | protein_coding |
| CENPA | 32 | protein_coding |

| | | |
|---|---|---|
| CIT | 32 | protein_coding |
| CKAP2L | 32 | protein_coding |
| GAS2L1 | 32 | protein_coding |
| HJURP | 32 | protein_coding |
| IQGAP3 | 32 | protein_coding |
| KIF2C | 32 | protein_coding |
| MTMR6 | 32 | protein_coding |
| NDC80 | 32 | protein_coding |
| NUSAP1 | 32 | protein_coding |
| PBK | 32 | protein_coding |
| RACGAP1 | 32 | protein_coding |
| RNU2.28P | 32 | snRNA |
| RNU6.1081P | 32 | snRNA |
| RNU6.760P | 32 | snRNA |
| SGOL2 | 32 | protein_coding |
| SKA1 | 32 | protein_coding |
| SKA3 | 32 | protein_coding |
| TACC3 | 32 | protein_coding |
| UBE2C | 32 | protein_coding |
| bP.2171C21.3 | 34 | NA |
| CDC45 | 34 | protein_coding |
| CDC6 | 34 | protein_coding |
| CLSPN | 34 | protein_coding |
| FAM161A | 34 | protein_coding |
| GINS2 | 34 | protein_coding |
| H2BFS | 34 | protein_coding |
| HIST1H2AD | 34 | protein_coding |
| HIST1H2AH | 34 | protein_coding |
| HIST1H2BJ | 34 | protein_coding |
| HIST1H2BK | 34 | protein_coding |
| HIST1H3A | 34 | protein_coding |
| HIST1H3B | 34 | protein_coding |
| HIST1H3H | 34 | protein_coding |
| HIST1H4C | 34 | protein_coding |
| HIST1H4D | 34 | protein_coding |
| LINC00505 | 34 | lincRNA |
| LINC00645 | 34 | lincRNA |
| LVCAT1 | 34 | lincRNA |
| MCM10 | 34 | protein_coding |

| | | |
|---|---|---|
| MCM2 | 34 | protein_coding |
| MCM3 | 34 | protein_coding |
| MCM6 | 34 | protein_coding |
| MIR4421 | 34 | miRNA |
| MIR874 | 34 | miRNA |
| MMS22L | 34 | protein_coding |
| MSH6 | 34 | protein_coding |
| NPAT | 34 | protein_coding |
| ORC1 | 34 | protein_coding |
| PCNA | 34 | protein_coding |
| RMI1 | 34 | protein_coding |
| RNU7.179P | 34 | snRNA |
| UG0898H09 | 34 | lincRNA |
| BLM | 38 | protein_coding |
| C16orf59 | 38 | protein_coding |
| CASP8AP2 | 38 | protein_coding |
| CDT1 | 38 | protein_coding |
| CEP85 | 38 | protein_coding |
| DKFZp779M0652 | 38 | lincRNA |
| DRAIC | 38 | lincRNA |
| DSCC1 | 38 | protein_coding |
| FANCC | 38 | protein_coding |
| FEN1 | 38 | protein_coding |
| HIST1H1C | 38 | protein_coding |
| HIST1H1E | 38 | protein_coding |
| HIST1H2BD | 38 | protein_coding |
| HIST1H2BO | 38 | protein_coding |
| HIST2H2AC | 38 | protein_coding |
| HIST2H2BF | 38 | protein_coding |
| HIST2H3A | 38 | protein_coding |
| HIST2H3C | 38 | protein_coding |
| HIST2H3D | 38 | protein_coding |
| HIST2H4A | 38 | protein_coding |
| HIST2H4B | 38 | protein_coding |
| RNU1.92P | 38 | snRNA |
| RNU6.48P | 38 | snRNA |
| RNU6.59P | 38 | snRNA |
| RP3.483K16.4 | 38 | NA |
| SUMO4 | 38 | protein_coding |

| | | |
|---|---|---|
| TICRR | 38 | protein_coding |
| XRCC2 | 38 | protein_coding |
| bP.2189O9.3 | 39 | NA |
| CTA.280A3.2 | 39 | NA |
| ERICD | 39 | lincRNA |
| FP671120.2 | 39 | NA |
| GS1.124K5.11 | 39 | NA |
| LINC00654 | 39 | lincRNA |
| LINC01091 | 39 | lincRNA |
| LINC01553 | 39 | lincRNA |
| MIR324 | 39 | miRNA |
| MIR5091 | 39 | miRNA |
| MIRLET7G | 39 | miRNA |
| RNU2.33P | 39 | snRNA |
| RNU2.38P | 39 | snRNA |
| RNU4.36P | 39 | snRNA |
| SNORA24 | 39 | snoRNA |
| SNORA60 | 39 | snoRNA |
| SNORA73B | 39 | snoRNA |
| CASC11 | 44 | lincRNA |
| CMB9.22P13.2 | 44 | NA |
| CPEB2.AS1 | 44 | lincRNA |
| DPH6.AS1 | 44 | lincRNA |
| DSCR9 | 44 | lincRNA |
| FLJ43879 | 44 | lincRNA |
| GS1.57L11.1 | 44 | NA |
| KIF25.AS1 | 44 | lincRNA |
| LINC00336 | 44 | lincRNA |
| LINC00421 | 44 | lincRNA |
| LINC00452 | 44 | lincRNA |
| LINC00460 | 44 | lincRNA |
| LINC00607 | 44 | lincRNA |
| LINC00705 | 44 | lincRNA |
| LINC00929 | 44 | lincRNA |
| LINC00940 | 44 | lincRNA |
| LINC01037 | 44 | lincRNA |
| LINC01085 | 44 | lincRNA |
| LINC01272 | 44 | lincRNA |
| LINC01320 | 44 | lincRNA |

| | | |
|---|---|---|
| LINC01337 | 44 | lincRNA |
| LINC01556 | 44 | lincRNA |
| LL0XNC01.7P3.1 | 44 | NA |
| MIR143HG | 44 | lincRNA |
| MIR340 | 44 | miRNA |
| MIR494 | 44 | miRNA |
| MIR532 | 44 | miRNA |
| MIR568 | 44 | miRNA |
| MIR6090 | 44 | miRNA |
| MIR769 | 44 | miRNA |
| RNU5A.4P | 44 | snRNA |
| RNU6.328P | 44 | snRNA |
| RNU6.409P | 44 | snRNA |
| RNU6.475P | 44 | snRNA |
| RP3.332B22.1 | 44 | NA |
| RP3.340N1.2 | 44 | NA |
| SFTA1P | 44 | lincRNA |
| SNORA1 | 44 | snoRNA |
| SNORA81 | 44 | snoRNA |
| SNORD37 | 44 | snoRNA |
| ZNF295.AS1 | 44 | lincRNA |
| C3orf35 | 46 | lincRNA |
| CARS2 | 46 | protein_coding |
| CTA.286B10.8 | 46 | NA |
| CTC.366B18.4 | 46 | NA |
| LINC00882 | 46 | lincRNA |
| LINC00964 | 46 | lincRNA |
| LINC01094 | 46 | lincRNA |
| MIR1275 | 46 | miRNA |
| MIR590 | 46 | miRNA |
| MIR6807 | 46 | miRNA |
| RASAL2.AS1 | 46 | lincRNA |
| RNU1.63P | 46 | snRNA |
| RNU2.48P | 46 | snRNA |
| RNU4.59P | 46 | snRNA |
| RNU6.1260P | 46 | snRNA |
| RNU6.353P | 46 | snRNA |
| RNU6.758P | 46 | snRNA |
| RP6.99M1.3 | 46 | NA |

| | | |
|---|---|---|
| SNORA75 | 46 | snoRNA |
| SUCLG2.AS1 | 46 | lincRNA |
| TEX41 | 46 | lincRNA |
| GTSE1.AS1 | 48 | lincRNA |
| LINC00235 | 48 | lincRNA |
| LINC00265 | 48 | lincRNA |
| LINC01315 | 48 | lincRNA |
| LINC01473 | 48 | lincRNA |
| LINC01541 | 48 | lincRNA |
| MIR3179.1 | 48 | lincRNA |
| MIR331 | 48 | miRNA |
| MIR3682 | 48 | miRNA |
| MIR425 | 48 | miRNA |
| MIR4522 | 48 | miRNA |
| RNU6.1229P | 48 | snRNA |
| RNU6.24P | 48 | snRNA |
| SNORA14B | 48 | snoRNA |
| SNORA70 | 48 | snoRNA |
| SNORA71B | 48 | snoRNA |
| SNORD38A | 48 | snoRNA |
| CCDC150 | 49 | protein_coding |
| CCDC28B | 49 | protein_coding |
| CCDC51 | 49 | protein_coding |
| CDC25A | 49 | protein_coding |
| DHFR | 49 | protein_coding |
| DNA2 | 49 | protein_coding |
| EXO1 | 49 | protein_coding |
| EZH2 | 49 | protein_coding |
| GMNN | 49 | protein_coding |
| HELLS | 49 | protein_coding |
| KLHL23 | 49 | protein_coding |
| LINC00383 | 49 | lincRNA |
| MIR16.2 | 49 | miRNA |
| MIR186 | 49 | miRNA |
| MIR188 | 49 | miRNA |
| MIR93 | 49 | miRNA |
| MIR941.2 | 49 | miRNA |
| ORC6 | 49 | protein_coding |
| PRIM1 | 49 | protein_coding |

| | | |
|---|---|---|
| SCARNA8 | 49 | scaRNA |
| SEPT7.AS1 | 49 | lincRNA |
| TIPIN | 49 | protein_coding |
| CCNB1 | 50 | protein_coding |
| CCNB2 | 50 | protein_coding |
| CDC20 | 50 | protein_coding |
| CDKN3 | 50 | protein_coding |
| DLGAP5 | 50 | protein_coding |
| ECT2 | 50 | protein_coding |
| FAM64A | 50 | protein_coding |
| HMMR | 50 | protein_coding |
| INCENP | 50 | protein_coding |
| KIAA0125 | 50 | lincRNA |
| KIF20A | 50 | protein_coding |
| KNSTRN | 50 | protein_coding |
| MIR33B | 50 | miRNA |
| NCAPD2 | 50 | protein_coding |
| PIF1 | 50 | protein_coding |
| RNU1.35P | 50 | snRNA |
| RNU6.1163P | 50 | snRNA |
| RNU7.3P | 50 | snRNA |
| TPX2 | 50 | protein_coding |
| UBALD2 | 50 | protein_coding |
| CDH1 | 52 | protein_coding |
| CYB561 | 52 | protein_coding |
| DDR1 | 52 | protein_coding |
| DUSP4 | 52 | protein_coding |
| ELF3 | 52 | protein_coding |
| FAM102B | 52 | protein_coding |
| FBP1 | 52 | protein_coding |
| FXYD3 | 52 | protein_coding |
| HEATR6 | 52 | protein_coding |
| IFI27 | 52 | protein_coding |
| IFI6 | 52 | protein_coding |
| ITPK1 | 52 | protein_coding |
| LINC01468 | 52 | lincRNA |
| METRN | 52 | protein_coding |
| MIR200B | 52 | miRNA |
| NR4A2 | 52 | protein_coding |

| | | |
|---|---|---|
| NRIP1 | 52 | protein_coding |
| PMEPA1 | 52 | protein_coding |
| PRLR | 52 | protein_coding |
| RAB25 | 52 | protein_coding |
| RNU6.813P | 52 | snRNA |
| RNVU1.19 | 52 | snRNA |
| SCARNA20 | 52 | scaRNA |
| SDC1 | 52 | protein_coding |
| SNORD104 | 52 | snoRNA |
| SSH3 | 52 | protein_coding |
| ST14 | 52 | protein_coding |
| SULT2B1 | 52 | protein_coding |
| TANC2 | 52 | protein_coding |
| TBX2 | 52 | protein_coding |
| TINCR | 52 | lincRNA |
| TMEM30B | 52 | protein_coding |
| TRIQK | 52 | protein_coding |

Supplementary table 3. Smart-seq-total primers used in the present study.

| | |
|---|---|
| Smart-seq-total oligo-dT | 5’-Biotin-CATAGTCTCGTGGGCTCGGAGATGTGTATAAGAGACAGT30VN-3’ |
| Smart-seq-total TSO | 5′-biotin-dUCGdUCGGCAGCGdUCAGdUdUGdUAdUCAACdUCAGACAdUrGrG+G-3′ |
| Smart-seq-total RV Amp | 5'-GTCTCGTGGGCTCGGAGATGTG-3' |
| Smart-seq-total FW Amp | 5'-TCGTCGGCAGCGTCAGTTGTATCAACT-3' |
| Smart-seq-total Custom SeqRead1 | 5’- TCGGCAGCGTCAGTTGTATCAACTCAGACATGGG-3’ |
